# Endothelial Cell Polarization and Orientation to Flow in a Novel Microfluidic Multimodal Shear Stress Generator

**DOI:** 10.1101/2020.07.10.197244

**Authors:** Utku M. Sonmez, Ya-Wen Cheng, Simon C. Watkins, Beth L. Roman, Lance A. Davidson

## Abstract

Endothelial cells (EC) respond to shear stress to maintain vascular homeostasis, and a disrupted response is associated with cardiovascular diseases. To understand how different shear stress modalities affect EC morphology and behavior, we developed a microfluidic device that concurrently generates three different levels of uniform wall shear stress (WSS) and six different WSS gradients (WSSG). In this device, human umbilical vein endothelial cells (HUVECs) exhibited a rapid and robust response to WSS, with the relative positioning of the Golgi and nucleus transitioning from non-polarized to polarized in a WSS magnitude- and gradient-independent manner. By contrast, polarized HUVECs oriented their Golgi and nucleus polarity to the flow vector in a WSS magnitude-dependent manner, with positive WSSG inhibiting and negative WSSG promoting this upstream orientation. Having validated this device, this chip can now be used to dissect the mechanisms underlying EC responses to different WSS modalities, including shear stress gradients, and to investigate the influence of flow on a diverse range of cells during development, homeostasis and disease.

## Introduction

Fluid shear stress, which arises from the intermolecular friction between differentially moving fluid layers, is an important hemodynamic factor that orchestrates the form and function of the blood vasculature through mechanosensitive signaling pathways. The inner lining of the vasculature is a single-layered squamous epithelium composed of endothelial cells (EC). Shear stress experienced by ECs at the vessel walls—known as wall shear stress (WSS)—has a profound influence on EC gene expression and behavior, with laminar shear stress inducing quiescence and disturbed (low or oscillatory) shear stress driving an inflammatory state ^1^. Additionally, laminar but not disturbed shear stress promotes EC elongation and cytoskeletal alignment parallel to the flow axis. Furthermore, in response to flow, ECs polarize, defined here as a distinct separation of the Golgi apparatus or microtubule organizing center (MTOC) and the nucleus. Polarized ECs generally orient against the direction of flow, with the Golgi or MTOC located upstream of the nucleus ^2-4^.

Although EC upstream orientation in response to flow is well documented, how this orientation affects EC function is not entirely clear. A lack of EC upstream orientation may be necessary but not sufficient for a pathological inflammatory response ^4^, and there is continuing debate regarding the correlation between Golgi-nucleus orientation and the direction of cell migration. Generally, in migrating single cells, the Golgi is positioned toward the leading edge, ahead of the nucleus ^5^. However, in ECs under flow, there are reports of correlation ^6,7^ and lack of correlation ^3,8,9^ between the orientation of the Golgi-nuclear vector and the direction of migration. Regardless, it is clear that Golgi-nucleus polarization and orientation are reproducible and sensitive EC responses to uniform laminar shear stress in vitro and in vivo ^4^.

There is also mounting evidence that ECs are not only sensitive to the WSS magnitude but also to the WSS gradient (WSSG) ^10-13^. During development, WSSGs help to shape the vasculature, with EC Golgi-nucleus orientation and migration toward regions of higher shear stress during vascular remodeling ^14,15^. Moreover, high WSSGs are associated with intracranial aneurysm development and atherosclerotic plaque instability ^16-19^. However, how WSSGs are sensed is poorly understood.

In vivo assays preserve the physiological complexity of the dynamic mechanical environment in the vasculature ^20^ but cannot decouple the role of shear stress from circulating factors. Furthermore, in vivo assays limit access to high-content imaging needed for mechanical characterization of blood flow and analysis of EC behavior. Moreover, natural variations in cardiovascular physiology limit reproducibility, hindering the establishment of precisely defined mechanistic relations between shear stress and EC biology. By contrast, microfluidic systems allow the generation of precisely controlled shear stress modalities in highly controlled microenvironments, and the effects of shear stress on pathologically-relevant EC phenotypes— such as cell alignment, polarization, intracellular organization, cytoskeletal rearrangement, apoptosis, and permeability—can be comprehensively investigated ^21^.

Despite the advantages of microfluidic devices, their full potential is rarely realized, as most systems fail to reproduce a broad spectrum of physiologically relevant shear stress magnitudes and gradients. Within the human blood vasculature, shear stress magnitudes range between 5 to 60 dyne.cm^−2 22^, and spatial shear stress gradients are predicted to be as high as 3 dyn.cm^−2^/μm ^23^. Yet, many microfluidic platforms limit experiments to a single shear stress condition and therefore require numerous repetitions with different flow rates to encompass physiologically-relevant shear stress modalities ^24-31^. Other microfluidic systems can simultaneously generate multiplexed shear stress regions within the same device ^32-38^. However, these systems are limited to a single mode of shear stress, with either shear stress gradient regions or uniform shear stress regions. Moreover, because these devices typically rely on parallel microchannels that branch from a common inlet channel via hydraulic resistance channels ^33,39^, they suffer from numerous limitations, including non-uniform cell seeding density that significantly affects the local flow fields ^40^, and localized flow perturbations affecting all microchannels ^41^. Additionally, each of the devices above require increased complexity of the flow circuit due to the need for an independent outlet for each perfusion channel.

To our knowledge, there is currently no microfluidic system that is capable of generating shear stress gradients in conjunction with different uniform shear stress regions for systematic analysis of shear stress-dependent cellular responses. Given that EC responses to flow may depend on the direction (i.e. sign), steepness (i.e., slope), and the average shear stress magnitude of the shear stress gradient ^11^, an ideal microfluidic system would allow simultaneous interrogation of EC behaviors in response to all these parameters. To address this need, we used 3D-printed molds to rapidly fabricate a novel microfluidic device that can simultaneously generate multiple shear stress modalities, allowing systematic mechanical screening and profiling of cells in physiologically-relevant dynamic flow conditions. Our microfluidic system is composed of microchannels with a constant height but variable widths, connected to each other in serial and parallel fashion. These microchannels comprise multiple shear stress regions, including three different magnitudes of uniform WSS and six different WSSGs (three positive, three negative). We characterized this system using analytical models and numerical simulations together with experimental validation.

To highlight the utility of this device, we quantitatively analyzed Golgi-nucleus relative position in human umbilical vein endothelial cell (HUVEC) monolayers using both endpoint and time lapse imaging. We found that flow-induced Golgi-nuclear polarization increased immediately after the onset of flow, concomitant with decreased Golgi and nuclear areas and increased Golgi-nuclear separation. However, while the frequency of polarized ECs was similar across all shear stress modalities, the frequency of ECs with Golgi-nucleus upstream orientation increased as WSS magnitude increased. Upstream orientation was also enhanced in regions of negative shear stress gradients and inhibited in regions of positive shear stress gradients.

## Results

### Overview of microfluidic device design

To study effects of different flow conditions on EC biology, we designed a microfluidic chip to meet several key performance and manufacturing criteria. (1) The chip allows generation of a wide range of WSS magnitudes and gradients. (2) The flow regimes are stable and robust such that external or internal disturbances do not interfere with the operation of the device. (3) The chip is compatible with high numerical aperture objectives for high content imaging. (4) Chip fabrication does not require the use of sophisticated equipment or specialized facilities.

To generate multiple flow regimes in a single device, our chip contains contiguous microchannel regions with different widths (Figure 1). The chip inlet leads to a narrow microchannel with constant width for high uniform WSS, which is followed by an expansion channel (Figure 1a; region 2) that generates a high-to-medium negative WSSG (i.e., WSS magnitude decreases in the direction of the flow). The next region is constant width for medium uniform WSS (Figure 1a; region 3), and is followed by an expansion channel (Figure 1a; region 4) that generates a medium-to-low negative WSSG. We designed these negative WSSG regions to allow comparison of different average WSS magnitudes within similar WSSGs. The medium-to-low WSSG region is followed by a constant-width region for low uniform WSS (Figure 1a; region 5), which then leads to two narrow constriction channels (Figure 1a; regions 6 and 7) that generate a very steep low-to high positive WSSG.

**Figure 1:**
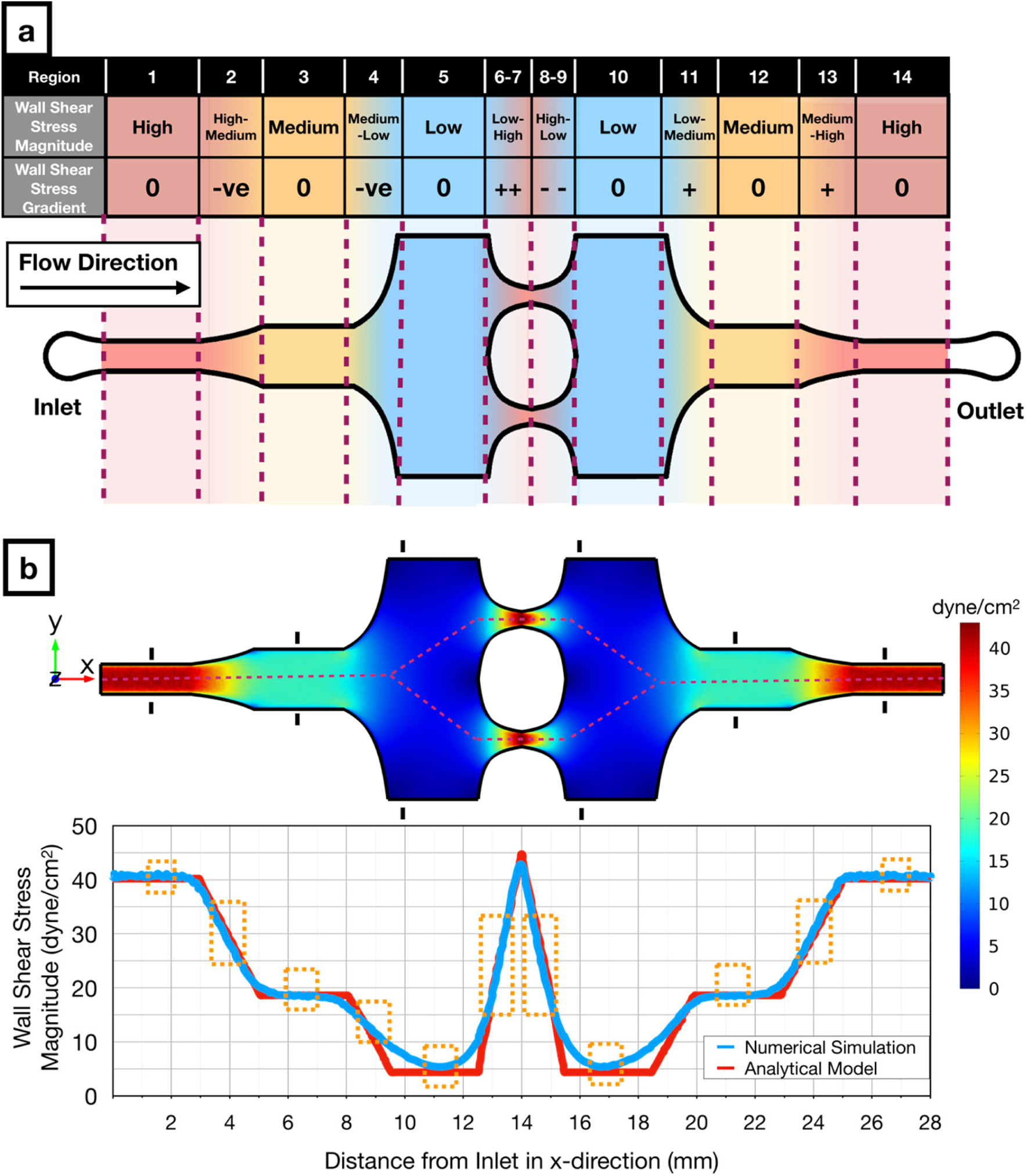
Design and numerical simulation of the novel microfluidic multimodal shear stress generator. a) Schematic of the microfluidic multimodal shear stress generator, indicating 14 regions with different shear stress modalities. b) Computational fluid dynamics simulations (Top) show the distribution of the shear stress across the microfluidic system. The black markers in each uniform shear stress region denote the hydraulic entrance length for those regions. Graph (Bottom) shows both the numerical (blue) and analytical (red) solutions for the magnitude of wall shear stress along the red dashed line (Top). Orange boxes (Bottom) indicate regions that offer predictable flow conditions where analytical and numerical solutions converge to similar results.

The microfluidic chip is symmetrical, with low (Figure 1a; region 10), medium (Figure 1a; region 12), and high (Figure 1a; region 14) uniform WSS regions in the outlet side of the microfluidic chip functioning as internal controls for their counterparts located on the inlet side of the chip. This duplication allows one to determine whether pressure drop across the microfluidic chip or factors secreted by cells in the upstream regions influences observed phenomena. Moreover, the microchannels with changing width in the outlet side of the chip (Figure 1a; regions 8, 9, 11 and 13) generate the same magnitude of WSSG as the microchannels in the inlet side of the chip (Figure 1a; regions 2, 4, 6, 7), but with the opposite sign. Altogether, our design enables a decoupling of effects of WSS magnitudes and gradients through 14 physiologically-relevant flow regions.

### Analytical and numerical characterizations of the microfluidic device

We designed and fabricated our microfluidic chip according to the following physical criteria. (1) The minimum microchannel width must be at least 200 µm since ECs auto-align in narrower microchannels in response to spatial confinement ^42^. (2) Uniform shear stress regions must be longer than hydrodynamic entrance length to ensure fully-developed flow (i.e., 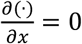). The hydrodynamic entrance length is calculated by

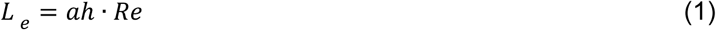

where *a* is an empirical proportionality constant equal to 0.08 for microchannels with rectangular cross-section, *h* is the height of the microchannel, and *Re* is the Reynolds number. (3) The thickness of the ceiling above the microchannels must be defined to minimize microchannel deformation under pressure-driven flow, based on published design suggestions ^43,44^. (4) Abrupt changes in the microchannel width must be avoided to prevent the generation of unwanted recirculation zones and vortices.

To characterize the effectiveness of our design, we performed both theoretical and computational analyses and measured flow velocity profiles in the microfluidic chip. In the microchannel regions with uniform width, the solution of the Navier-Stokes equations for the steady-state laminar flow approximates to Hele-Shaw flow, where the WSS acting on the cells can be calculated as

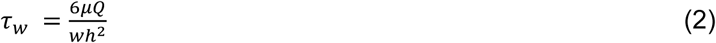

where *μ* is the dynamic viscosity of the fluid, *Q* is the volumetric flow rate, and *w* is the width of the microchannel. However, especially in high WSS regions where the aspect ratio (*h*/*w*) can be as high as 0.2, the sidewalls also contribute to the WSS acting upon the cells. This sidewall effect can be taken into account by using the empirically-corrected version of the WSS formula above, which is valid for aspect ratios up to 0.3 ^45^. This corrected formula is

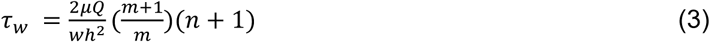

where *n =* 2 and *m* is defined as

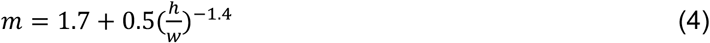

On the other hand, microchannel regions with changing width must have linear streamwise (i.e., along the *x* axis) WSSGs so that cells in a given region are exposed to the same gradient slope. The WSS formula for the WSSG can be written as

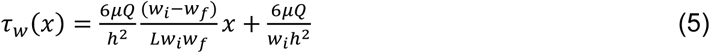

where *w*_*i*_ and *w*_*f*_ are the width of the microchannel at the beginning and the end of the WSSG region, respectively, while *L* is the axial length of the WSSG region (Supplementary Figure 1). The curved microchannel sidewall formula needed to obtain this WSSG can also be written as

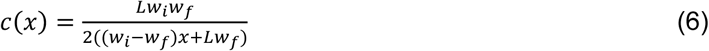

Using these formulae, we generated the microchannel sidewall profiles with respect to the initial and final microchannel widths for the desired WSSG slope. (See Supplementary Information for the derivation of these formulae).

Dynamic viscosity (µ) in the WSS formulae above is a material property and its value is specific to the perfusion medium and its temperature. However, in the literature, it is common to assume the dynamic viscosity of the perfusion medium is equal to that of water at room temperature ^35,46^, whereas other studies have assumed a much higher viscosity ^47^. To ensure our WSS calculations were correct, we measured the dynamic viscosity of the complete EC growth media that we used in this study and found its value as 0.87 mPa.s at 37 °C (Supplementary Figure 2).

Using the analytical model and the measured dynamic viscosity, we defined the exact dimensions of each microchannel region (Supplementary Figure 3) which resulted in uniform WSS magnitudes of 40.2 dyne.cm^−2^, 18.6 dyne.cm^−2^, and 4.4 dyne.cm^−2^ for high (regions 1 and 14), medium (regions 2 and 13), and low (regions 5 and 10) uniform WSS regions, respectively, at the flow rate of 1.6 ml.min^−1^. Slopes of the linear WSSGs between these regions had absolute magnitudes of 8.5 dyne.cm^−2^.mm^−1^ (regions 2,4,11,13) and 17.4 dyne.cm^−2^.mm^−1^ (regions 6,7,8,9). Numerical simulations yielded similar results across the microfluidic chip (Figure 1b) with 659 Pa pressure drop across the entire microchannel. Numerical simulations also revealed how far the wall effects ^48^ propagated into the middle of the microchannel from the sidewalls. For subsequent cellular analysis, we only included the microchannel portions sufficiently far from to the sidewalls where the consistent shear stress values were predicted from analytical flow analysis (Supplementary Figure 4). Lastly, we marked the hydrodynamic entrance length for each uniform shear stress region (Figure 1b; black markings) and did not analyze the cells in microchannel regions with potentially undeveloped flow. These data demonstrate that our microfluidic system design can span a 10-fold wall shear stress range and the spatial gradients within this range, covering a majority of physiologically-relevant flow modalities observed in vascular networks^49^.

### Fabrication and experimental validation of the microfluidic device

Our design flexibly connects all shear stress regions in a single layer with only one inlet and one outlet, facilitating fabrication, operation, and imaging. Using stereolithographic additive manufacturing technology ^50^, we fabricated the molds for our microfluidic device in less than 4 hours in a fully-automated manner without the need for cleanroom facilities. We used these molds to rapidly prototype the PDMS (polydimethylsiloxane)-based microfluidic chips that we used throughout this study (Figure 2a). However, additive manufacturing can introduce higher levels of surface roughness than photolithographic microfabrication, which might result in fluctuations in WSS. To assess surface roughness, we imaged the microchannels using scanning electron microscopy (SEM). Images revealed surface roughness only at a submicron level (Figure 2b), which is less than 0.5% of the microchannel height, introducing variation that would not likely impact the flow perceived by cultured cells. In support of this assumption, micro-particle image velocimetry (µPIV) measurements with fluorescent beads showed parabolic velocity profiles across the height of the microchannels (Figure 2c), as predicted by the Hele-Shaw flow model. Therefore, we conclude that stereolithography is a valid rapid fabrication technique able to meet the geometrical requirements of our microfluidic chip designs.

**Figure 2:**
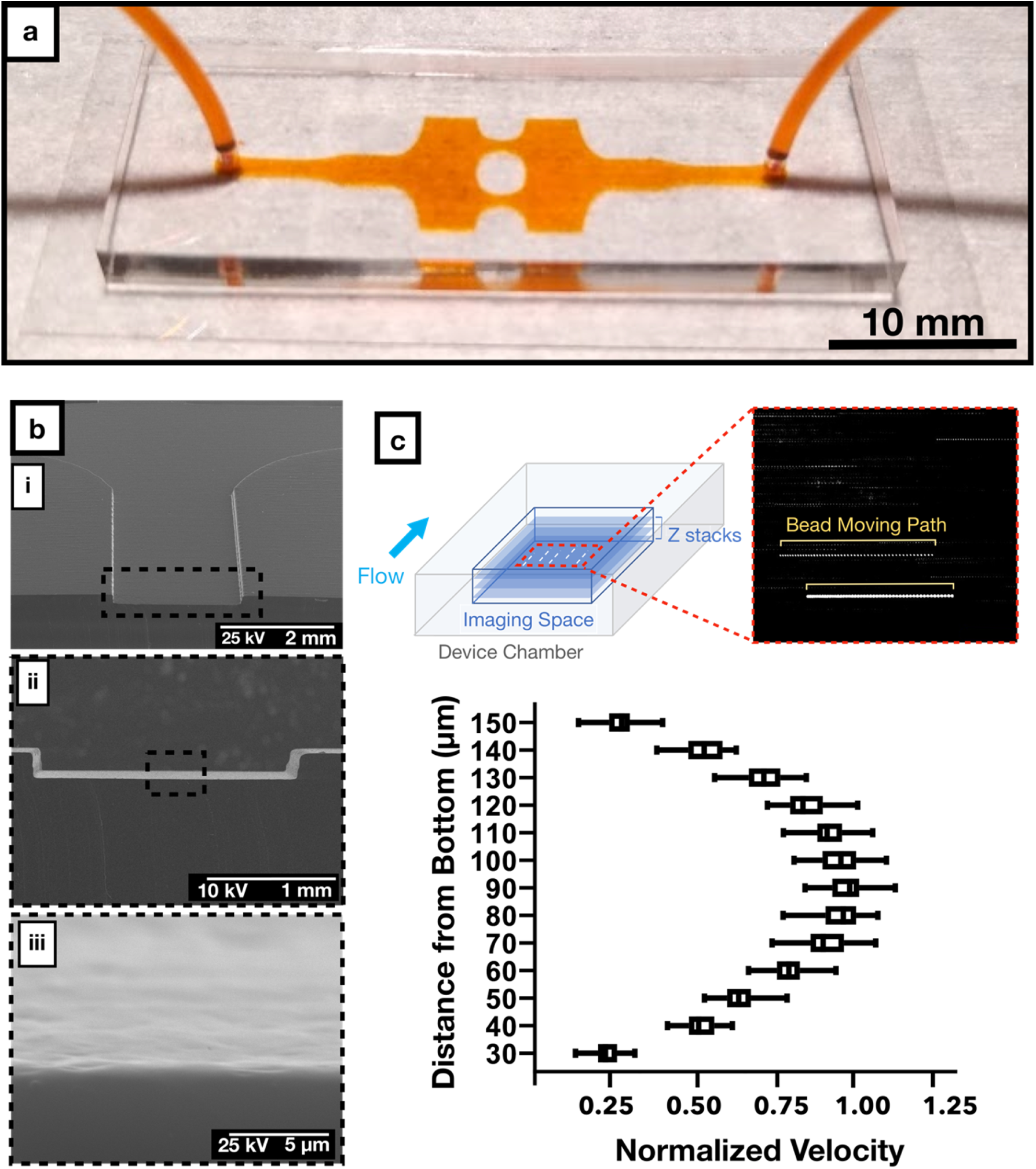
Fabrication and experimental characterization of the microfluidic chip. a) Representative sample of our PDMS-glass microfluidic chip with the inlet and outlet tubing connected. Microchannels were filled with orange dye to facilitate visualization. b) Scanning electron micrograph of the PDMS microchannel with increasing magnification, displaying the surface roughness of the microchannel. c) Flow profile at the cross section of the microfluidic device plotted from micro-particle image velocimetry analysis at 0.5 ml/min. Each dashed line in the inset represents the path for each fluorescence bead in the field.

We designed our microfluidic chip to ensure laminar flow, free from flow separation and consequent vortices as well as inertial drag forces. To confirm laminar flow conditions within the microfluidic chip, we used fluorescent beads in the perfusion medium to experimentally visualize the streaklines throughout the microchannels. In these experiments, we observed no flow recirculation or vortex generation, validating our flow model (Supplementary Movie 1). Furthermore, beads were randomly distributed across the channel width even in high WSS regions, confirming the lack of inertial drag forces. These observations confirm that flow regimes within our device maintain smooth WSS transitions with predictable WSSGs. Altogether, these characterization studies show that our novel microfluidic chip can stably generate different physiologically-relevant shear stress modalities in a single device, as predicted by our design calculations.

### Perfusion set-up and automated analysis of Golgi and nucleus

To assess effects of WSS on EC biology, we created a sterile flow circuit in which a peristaltic pump drew medium from a heated reservoir and passed it through a pulse dampener and bubble filter before entering the microfluidic chip (Figure 3a). After seeding HUVECs into the chip, we cultured them in static conditions for the establishment a uniform monolayer across the chip (Supplementary Figure 5) prior to application of different modalities of physiological flows for 24 hours (Figure 3b). We then fixed and stained cells for Golgi and nucleus. For all morphological analyses, we only imaged regions of the microfluidic channel where simulation and analytic solutions maintained concordance and where the cells were buffered from gradients due to wall effects. From these images, we quantitatively characterized HUVEC polarity responses to flow using a custom-written macro. This macro (Supplementary Figure 6) automatically paired nuclei and Golgi and measured Golgi-nucleus polarization and orientation of thousands of cells (Figure 3c) for subsequent statistical analysis.

**Figure 3:**
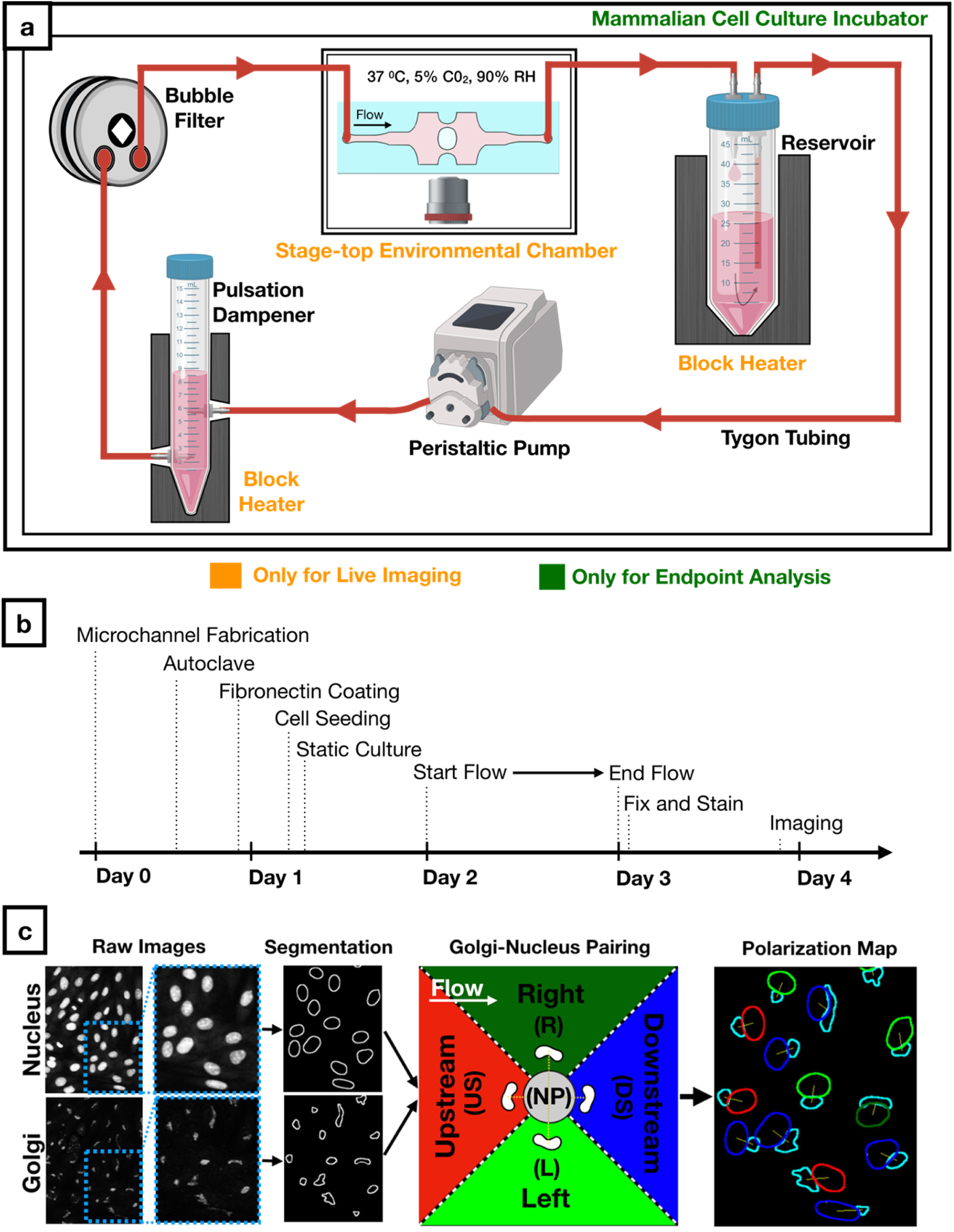
Workflow for cell culture experiments and image analysis. a) Schematic of the sterile flow circuit used with the microfluidic chip. The flow circuit can be set up on a microscope stage for live imaging under flow or it can be placed in a cell culture incubator for endpoint analysis. b) Experimental timeline for Golgi-nucleus morphology studies. Human umbilical vein endothelial cells (HUVECs) were seeded inside the microfluidic chip 16 hours before initiating flow and fixed after 24 hours of flow. c) Fluorescent images of nuclei and Golgi were analyzed automatically using a custom-written macro to define and pair each nucleus and Golgi as well as to categorize each cell as nonpolarized (gray) or polarized. Polarized cells were binned into one of four Golgi-nucleus orientation categories: upstream (red), downstream (blue), right (dark green), left (light green), with respect to the direction of flow. Orientation maps outline Golgi (cyan) and nucleus, with nucleus color indicating the orientation of the cell.

### Golgi-nucleus polarization is more sensitive than Golgi-nucleus orientation to uniform WSS

The separation of the Golgi and nucleus in the plane of the EC sheet (i.e., Golgi-nucleus polarization) and their alignment with respect to the direction of fluid flow (e.g., Golgi-nucleus orientation) are robust responses to shear stress ^4^. In our microfluidic chip under static conditions, approximately 50% of HUVECs were unpolarized and 50% were polarized but not oriented towards a particular direction (Figure 4a, b-i). All microchannel regions showed similar results in static conditions, indicating that the microchannel geometries did not alter either the polarization or the orientation of ECs (see Supplementary Table 1a for the complete list of exact p-values). When we exposed HUVECs to laminar flow for 24 hours, approximately 95% of cells became polarized in all uniform WSS regions, regardless of the WSS magnitude (Figure 4a, b-ii). Similar to the static conditions, HUVECs oriented randomly in response to low uniform WSS (4 dyne.cm^−2^). By contrast, HUVECs oriented upstream in a shear stress magnitude dependent manner at 19 and 40 dyne.cm^−2^ (see Supplementary Table 1b for the complete list of exact p-values). We obtained similar results from both inlet and outlet sides, indicating that the pressure drop across the microchannels did not influence Golgi-nucleus polarization or orientation. These results suggested that even at 4 dyne.cm^−2^, which is considered to be the very lowest end of the physiological shear stress range for vascular ECs ^51^, WSS was sufficient to induce Golgi-nucleus polarization (see Supplementary Table 1c for the complete list of exact p-values) but not Golgi-nucleus upstream orientation.

**Figure 4:**
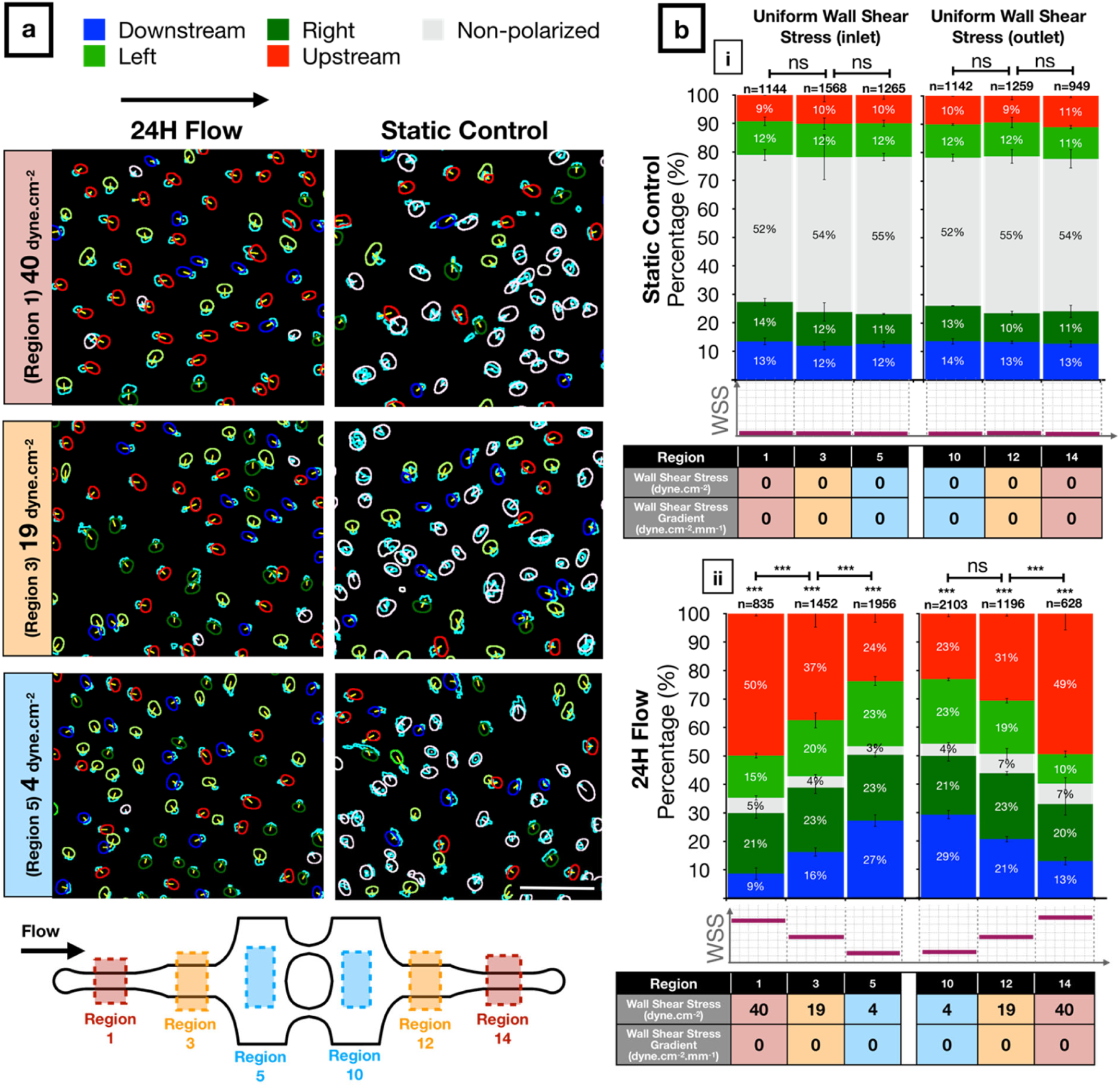
Golgi-nucleus polarization and orientation in response to uniform wall shear stress (WSS). a) Golgi-nucleus polarization and orientation maps from a representative 24-hour experiment, showing HUVECs exposed to high (40 dyne.cm^−2^; region 1), medium (19 dyne.cm^−2^; region 3), and low (4 dyne.cm^−2^; region 5) uniform WSS magnitudes, together with the static controls from the respective microchannel regions. Golgi are outlined in cyan. To indicate Golgi-nucleus orientation, nuclei are outlined in red (upstream), blue (downstream), dark green (right), light green (left), or white (nonpolarized). Scale bar, 100 µm. b) Quantitative analysis of the polarization and orientation responses in (i) static control and (ii) uniform WSS of indicated magnitudes (table below bar graphs), with colors corresponding to orientation category. n = number of cells analyzed in each region, combined over 3 independent experiments. Data are mean ± SEM. Chi-square independence test followed by post-hoc adjusted residual test with Bonferroni correction. ***p<0.0005; ns, not significant at p > 0.01 after Bonferroni correction. See Supplementary Table 1 for the complete list of exact p-values.

### Negative WSSGs promote and positive WSSGs inhibit Golgi-nucleus upstream orientation

Next, we assessed Golgi-nucleus orientation in WSSG regions. In these regions, the percentage of polarized cells increased from ∼50% in static controls to ∼95% with flow (Figure 5), similar to results in uniform WSS regions. However, the Golgi-nucleus orientation differed based on the sign and the steepness of the WSSG. More HUVECs were oriented upstream in response to a negative WSSG compared to a positive WSSG of the same slope and average WSS magnitude (Figure 5b; see Supplementary Table 1b for the complete list of exact p-values). The difference between numbers of upstream and downstream oriented ECs increased as the negative WSSG steepened (Figure 5b-iii, iv). For example, in response to a steep negative WSSG (−17 dyn.cm^−2^.mm^−1^) with average WSS magnitude of 24.5 dyn.cm^−2^, cells oriented upstream similarly to cells exposed to uniform WSS magnitude of 40 dyn.cm^−2^ (Figure 5b, Regions 8 & 9 vs. Figure 4c, Regions 1 & 14). By contrast, positive WSSG had the opposite effect, reducing numbers of upstream oriented ECs. For example, in response to a shallow positive WSSG (8 dyn.cm^−2^.mm^−1^) with average WSS magnitude of 11.5 dyn.cm^−2^, cells oriented randomly, similarly to cells in uniform WSS magnitude of 4 dyn.cm-2. (Figure 5b, Region 11 vs. Figure 4b-ii, Regions 5 & 10; see Supplementary Table 2 for the complete list of exact p-values).

**Figure 5:**
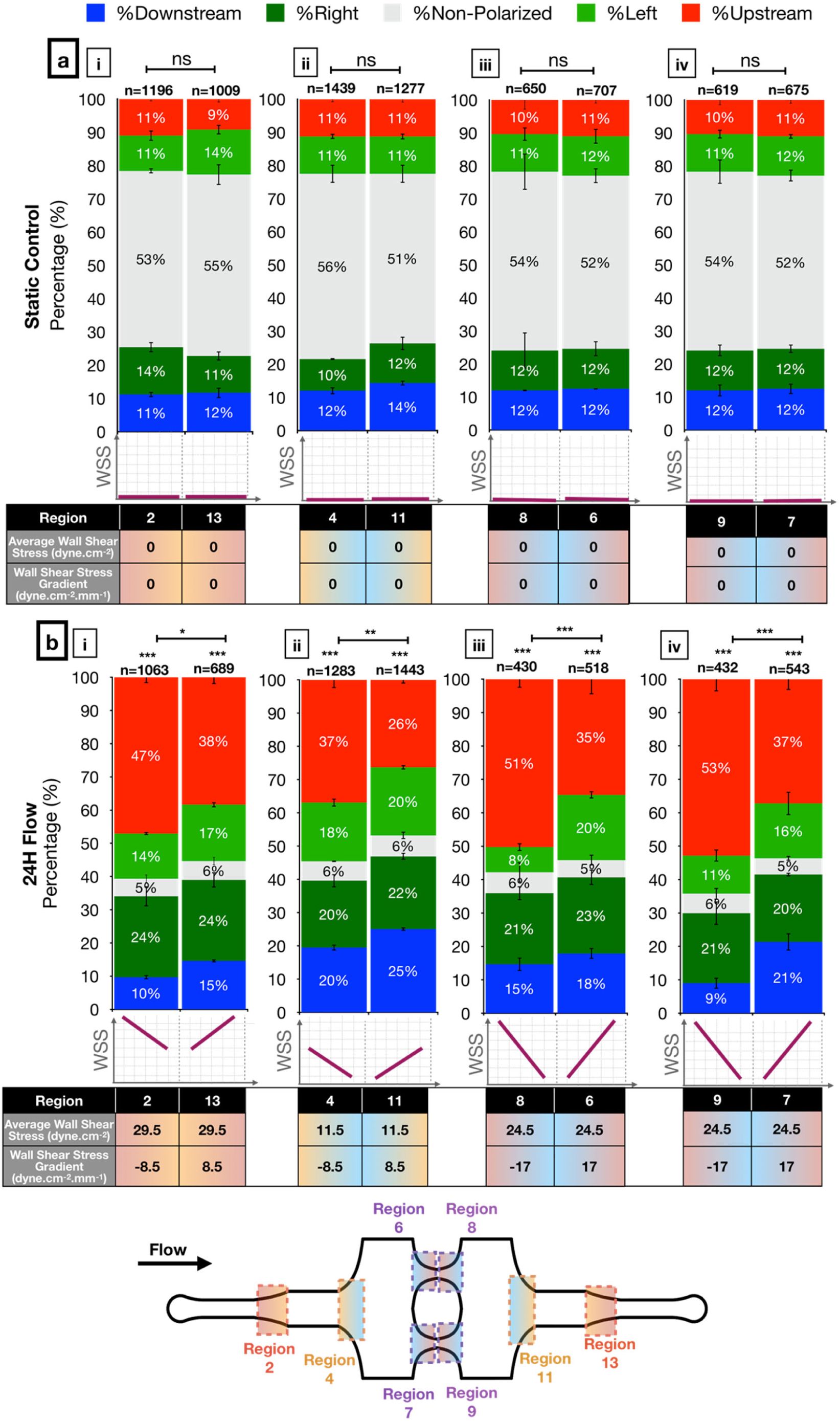
Golgi-nucleus polarization and orientation in response to wall shear stress gradients (WSSG). Quantitative analysis of the polarization and orientation responses in (a) static control and (b) WSSGs, comparing regions of oppositely orientated WSSGs of low steepness and high average WSS (b-i), low steepness and low average WSS (b-ii), and high steepness and medium average WSS (b-iii, iv), with average WSS and WSSG as indicated the table below the bar graphs. Golgi-nucleus orientation is indicated as upstream (red), downstream (blue), right (dark green), left (light green), or nonpolarized (gray). n = number of cells analyzed in each region, combined over 3 independent experiments. Data are mean ± SEM. Chi-square independence test followed by post-hoc adjusted residual test with Bonferroni correction. ***p<0.0005; **p<0.005; *p<0.01; ns, not significant at p > 0.01. See Supplementary Table 1 for the complete list of exact p-values.

To confirm that our observations regarding effects of WSS and WSSGs did not depend on the specific geometry of our microfluidic chip, we modified the width and the length of the device following the design guidelines described above to generate higher WSS magnitudes with steeper WSSGs. HUVECs exposed to WSS magnitudes of up to 60 dyne.cm^−2^ and gradients up to 40 dyne.cm^−2^.mm^−1^ showed very similar polarization and orientation to cells in microchannels with standard geometry (Supplementary Figure 7). Thus, the effects of WSS and WSSG on Golgi-nucleus polarization and orientation did not depend on the specific microfluidic device design.

### Golgi-nucleus upstream orientation develops quickly after the onset of flow

To understand the dynamics of EC responses to WSS, we live-imaged HUVECs in our microfluidic device, with a similar set-up to that described above, but with the microfluidic chip housed in a stage-top environmental chamber (Figure 3a). We used Hoescht 3342 dye to label nuclei and baculoviral-transfected GFP-tagged N-acetylgalactosaminyltransferase to label Golgi, and imaged across the entire chamber at 15 minute intervals over a 6 hour period. In the static controls, approximately half of the cells were unpolarized while the rest were randomly oriented at all time points (Figure 6a). Although individual cells were constantly changing their state of polarization and direction of orientation, the population level response remained the same over time (Figure 6a and Supplementary Movie 2). Upon exposure to flow, the number of upstream oriented cells increased while the number of downstream oriented cells decreased (Figure 6a, Supplementary Movie 3). The ratio of upstream oriented cells to downstream oriented cells (orientation index) increased over time, with the greatest rate of change in the first 3 hours, regardless of shear stress modality (Figure 6b). The orientation index increased more rapidly in response to higher WSS magnitude (Figure 6b-i), and negative WSSGs resulted in higher orientation index compared to positive WSSGs with the same average WSS magnitude (Figure 6b-ii, iii). Overall, results from live imaging experiments supported observations of the Golgi-nucleus polarization and orientation in response to different physiological WSS modalities after 24 hours of flow and demonstrated that the Golgi-nucleus polarization and orientation response develops quickly after the onset of the flow.

**Figure 6:**
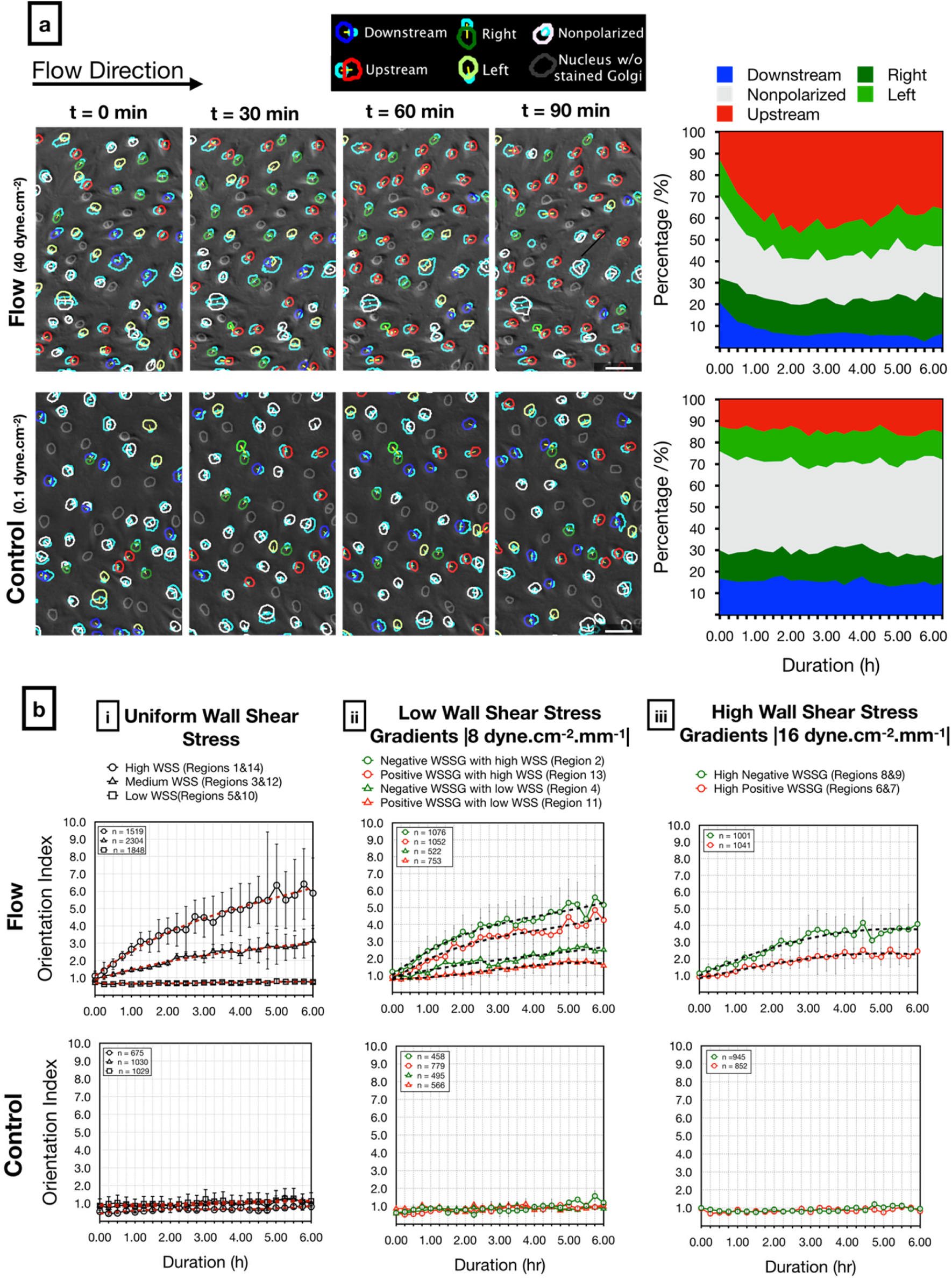
Live analysis of Golgi-nucleus polarity and orientation in response to shear stress. a) Representative images (left) from a time-lapse series of live-stained HUVECs (Golgi, nucleus) exposed to high uniform WSS (top; 40 dyne.cm^−2^) or quasi-static conditions (bottom; 0.1 dyne.cm^−2^). Overlays show Golgi-nucleus orientation maps merged with DIC images. Golgi are outlined in light blue. To indicate Golgi-nucleus orientation, nuclei are outlined in red (upstream), blue (downstream), dark green (right), light green (left), white (nonpolarized), or gray (Golgi not visible). Stacked area plots (right) demonstrate dynamic Golgi-nuclear orientation map over 6 hours of live imaging. Scale bars, 50 µm. b) Orientation index (ratio of upstream oriented cells to downstream oriented cells) of HUVECs exposed to flow (Top) or static (Bottom) conditions over 6 hours, in uniform WSS regions (i), low WSSG regions (ii), and high WSSG regions (iii). n = number of cells analyzed in each group, combined over 4 independent experiments for flow and 3 independent experiments for control. Data are mean ± SEM.

### ECs respond to flow by altering Golgi and nucleus sizes and Golgi-nucleus distance but these changes are independent of WSS magnitude or gradient

Although we found that the Golgi-nucleus orientation was regulated by WSS magnitude and gradient, the flow-dependent transition from a nonpolarized state to a polarized state did not depend on WSS magnitude or gradient. To investigate EC morphological response to flow, we measured the projected area of the nucleus and Golgi, and the distance between their centers after 24 hours of flow (Figure 7). We found that the nucleus and Golgi areas were significantly decreased under flow by 20% and 51%, respectively, compared to static controls (Figure 7a, b). On the other hand, the average distance between the Golgi and nucleus centers significantly increased by 34%, from 6.6 µm in static conditions to 8.9 µm under flow (Figure 7c; see Supplementary Table 3a for the complete list of exact p-values). All of these changes were independent of WSS magnitude and gradient (see Supplementary Table 3b for the complete list of exact p-values).

**Figure 7:**
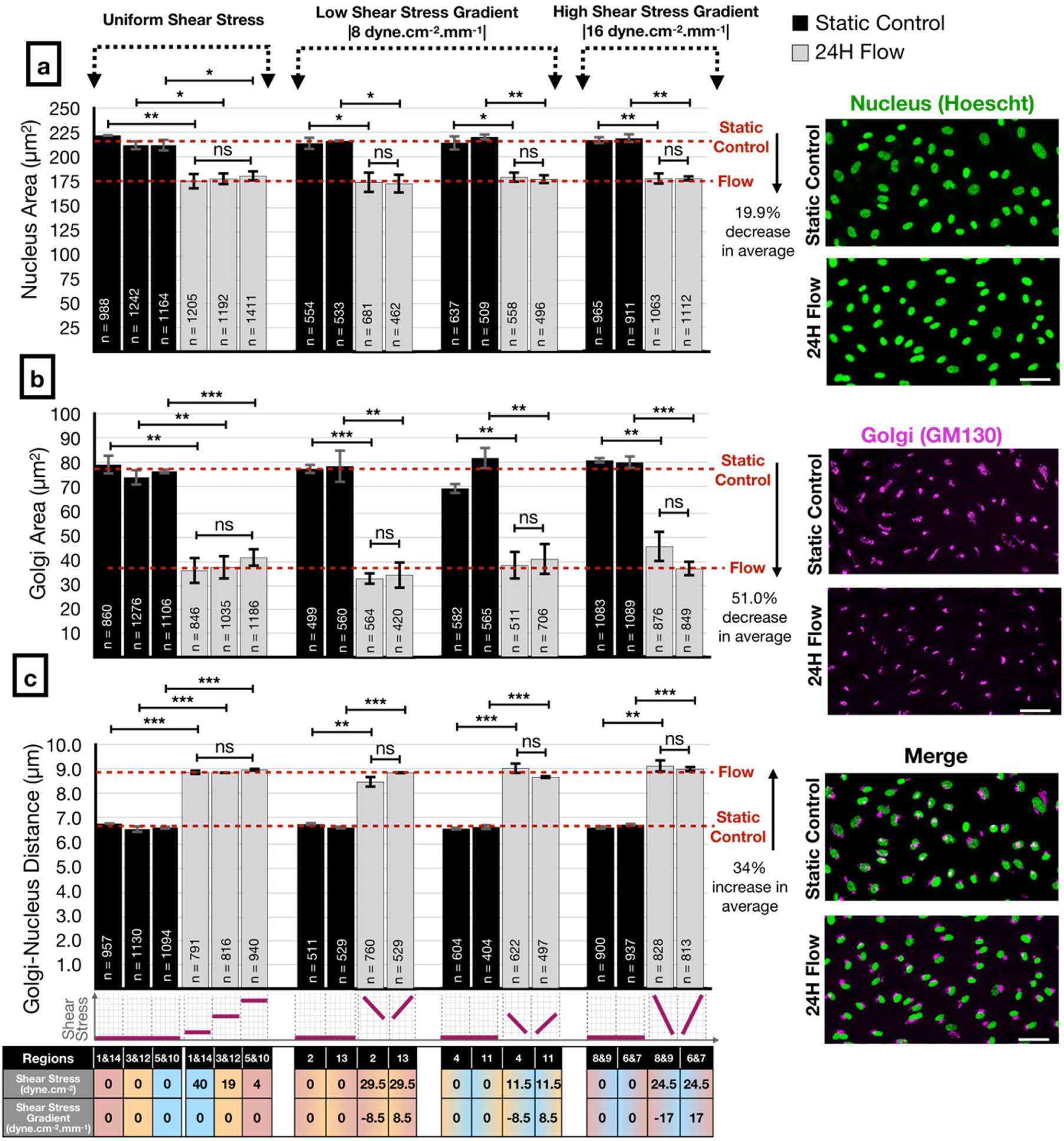
Analysis of Golgi and nucleus sizes and the distance between them in response to different modalities of shear stress. Human umbilical vein endothelial cell (HUVEC) nuclear projected area (a), Golgi projected area (b), and Golgi-nucleus distance (c) assayed after 24 hours of static (black bars) or flow (gray bars) conditions, in uniform WSS regions (left), low WSSG regions (center), and high WSSG regions (right) of the microfluidic device. Representative images of Golgi stain (GM130) and nuclear stain (Hoescht 33342) at far right are from region 1. Scale bars, 50 µm. n = number of cells analyzed in each group, combined over 3 independent experiments. Data are presented as mean ± SEM. ANOVA followed by post-hoc two-tailed unpaired student’s t-test with Bonferroni correction for pairwise comparison. ***p<0.001; **p<0.01; *p<0.05; ns, not significant at p > 0.05. See Supplementary Table 3 for the complete list of exact p-values.

To evaluate the dynamics of observed increases in Golgi-nucleus distance, we measured the change in this distance over 6-hours from time lapse sequences (Figure 8a). Golgi-nucleus distance increased immediately after the onset of flow, reached a peak value within 3 hours, then remained stable and at a similar level regardless of WSS modality (Figure 8b). By contrast, in the static control experiments, the Golgi-nucleus distance did not increase, although distances fluctuated over time. Together, these data suggest that Golgi-nucleus polarization is an early and uniform response to WSS. Both the decrease in the Golgi and nucleus areas as well as the increase in the distance between them contribute to polarization by decreasing their overlap and rendering them as two distinct entities. On the other hand, Golgi-nucleus upstream orientation in response to flow occurs within a similar timeframe in all tested flow modalities, but the extent of this response depends on the magnitude of the WSS and the sign of the WSSG.

**Figure 8:**
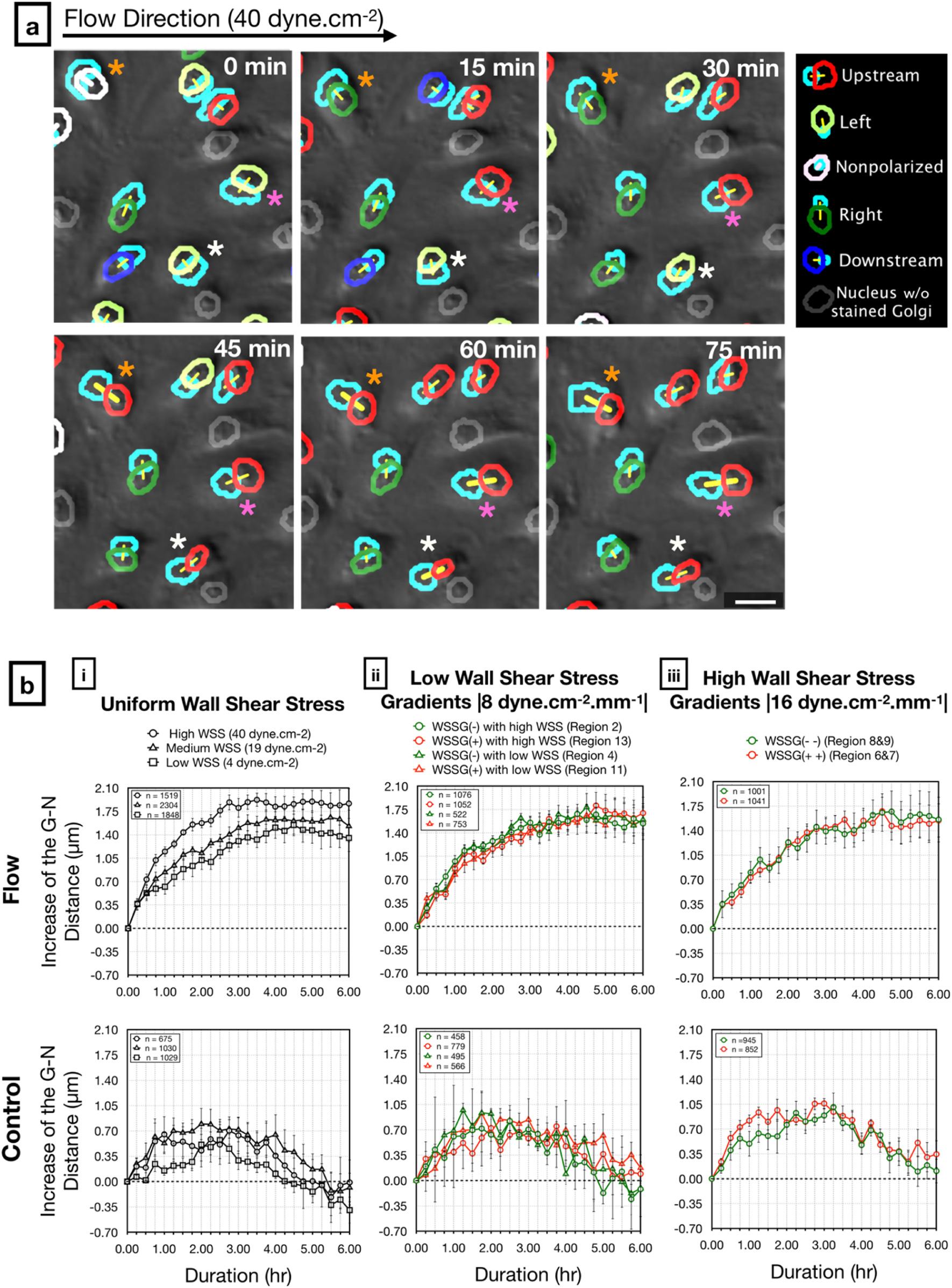
Live analysis of Golgi-nucleus distance in response to shear stress. a) Representative images from a time-lapse series of live-stained HUVECs (Golgi, nucleus) exposed to high uniform WSS (40 dyne.cm^−2^). Overlays show Golgi-nucleus orientation maps merged with DIC images. Golgi are outlined in light blue. To indicate Golgi-nucleus orientation, nuclei are outlined in red (upstream), blue (downstream), dark green (right), light green (left), white (nonpolarized), or gray (Golgi not visible). The centers of paired Golgi and nuclei are connected with a yellow line. Differently colored asterisks mark individual cells demonstrating increasing Golgi-nucleus distance over time. Scale bar, 25 µm. b) Quantitative analysis of the Golgi-nucleus (G-N) distance in HUVECs exposed to flow (top) or static (bottom) conditions over 6 hours, in uniform WSS regions (i), low WSSG regions (ii), and high WSSG regions (iii). n = number of cells analyzed in each group, combined over 4 independent experiments for flow and 3 independent experiments for control. Data are mean ± SEM.

## Discussion

Our microfluidic chip allows interrogation of effects of a wide range of physiologically-relevant shear stress magnitudes and shear stress gradients on EC morphology and behavior in a single experiment. The design strategy generates three different magnitudes of uniform shear stress and six different spatial shear stress gradients in a single unit with a closed-loop flow circuit that has only one inlet and one outlet. With the experimental flow rate of 1.6 ml.min^−1^, we achieved laminar flow; the maximum Reynolds number inside the chip was ∼51, which was more than an order of magnitude lower than the threshold value of ∼2300 for turbulent transition. Furthermore, characterization of this chip demonstrated flow stability and validated the analytical and numerical models used to design the system. Importantly, our chip allowed application of a wide range of shear stress without progressive deviation from target magnitudes, demonstrating improved performance compared to other designs ^38^.

For most microfabrication techniques, fine control of microchannel width is easier than fine control of microchannel height. Moreover, because WSS scales with *h*^−2^ and *w*^−1^, small deviations in microchannel height cause more pronounced deviations in shear stress than small deviations in channel width. Therefore, using varying microchannel widths to generate multiple shear stress modalities both simplifies microfabrication and improves accuracy.

The design of our microfluidic system allowed for long-term cell culture and minimal reagent use. The 200 µm microchannel height kept HUVECs viable for 24 hours in static culture while requiring only ∼30 mL perfusion media for application of shear stress. Moreover, the microchannel height was considerably larger than the thickness of the EC monolayer ^52^, allowing us to ignore effects of dynamic EC height changes in response to shear stress ^53^.

One of the biggest obstacles to widespread adoption of “lab-on-a-chip” technologies is the complexity of integrating microfluidic chips with bulky fluid handling systems, e.g. the “lab-around-the-chip.” ^54^. Such obstacles can be removed by miniaturization and on-chip integration of the equipment required to run the microfluidic systems. To this end, we propose future designs that integrate a continuous bubble filter ^55^ using the multilayer microfabrication capability of stereolithography-based 3D printing (see addition of reservoir and discharge channel, Supplementary Figure 8). Further integration of bulk fluid handling devices with the chip could render this microfluidic system even more accessible to most laboratories.

Results from our microfluidic device are concordant with published findings that ECs adopt upstream orientation after 3 hours exposure to shear stresses above 7.2 dyne.cm^−2 2^. With a more detailed analysis of cell polarity and orientation, we found that shear stress as low as 4 dynes.cm^−2^ led to a rapid and dramatic increase in numbers of polarized HUVECs compared to static controls, independent of shear stress magnitude. Analysis of nuclear and Golgi size and position suggested that this increase was likely due to both nuclear and Golgi compaction combined with increased Golgi-nucleus distance, which were similar across all shear stress modalities. Based on reports of compact organellar morphologies in G1-phase hTert-RPE1 cells ^56^ and reports that flow leads to cell cycle arrest at G1/S transition in HUVECs ^57^, we speculate that increased polarization reflects flow-mediated cell cycle arrest. However, our current findings are purely correlative; we do not know whether cell cycle arrest is necessary or sufficient for the Golgi-nuclear changes observed here. Future work with our device would allow a more detailed dissection of these processes.

The effect of shear stress gradients on EC morphology and function is not well understood. ECs surrounding a stenosis or atherosclerotic plaque, which are subjected to spatial WSSGs, adopt a cobblestone as opposed to an elongated morphology ^58^, and associated EC dysfunction may contribute to plaque rupture ^59^. The inhibition of EC alignment, e.g. cell shape change, with positive WSSG and its promotion with negative WSSG have been demonstrated in macroscale flow chambers and T-shaped perfusion channels at supraphysiological shear stress levels ^60,61^. Here, we extend these observations on whole cell morphology, demonstrating that a physiologically-relevant positive WSSG inhibits Golgi-nucleus upstream orientation whereas a negative WSSG enhances upstream orientation. Whether these changes in Golgi-nuclear polarization and orientation correlate with changes in directional migration or EC function remain to be determined.

Whether Golgi-nucleus polarization and orientation reflect simple mechanical responses to flow, e.g. displacement of the nucleus under a load, as opposed to mechanotransduction-mediated responses, remains unclear. The mechanical hypothesis suggests that the nucleus is dragged by the flow to downstream regions of the cytoplasm ^2^. Additionally, the tensile and compressive forces generated along the direction of the flow by the positive and negative WSSGs, respectively, may contribute to the extent of upstream orientation ^10,17^. Together, our results suggest that polarization and orientation may be governed by different mechanisms. For instance, polarization may result from mechanotransduction leading to flow-dependent cell cycle arrest, whereas upstream orientation may result from integrating both the physical and biochemical responses to shear stress magnitude and gradient. Further research will be required to systematically test the roles of different flow modalities in triggering mechanosensitive cell polarity signaling pathways such as NOTCH1 ^62^, PAR-3 ^4^, and APLNR ^6^ in shear stress-mediated Golgi-nucleus polarization and orientation.

## Materials & Methods

### Microchannel fabrication

Microfluidic channels were fabricated out of Polydimethylsiloxane (Sylgard 184 PDMS; Dow Corning, USA) using replica molding technique with 3D-printed molds. First, negatives of the microchannel geometries were modelled in SolidWorks 2016 (Dassault Systèmes, Vélizy-Villacoublay, France). The molds were manufactured using VIPER si2T Stereolithography System (3D Systems, Rock Hill, South Carolina, USA) in high quality mode with Accura SI 10 Polymer as the photocurable resin. Fabricated molds were post-cured with UV and their functional surfaces were coated with Tridecafluoro-1,1,2,2-tetrahydrooctyl-1-trichlorosilane (Cat: T2492, United Chemical Technologies, USA). PDMS was mixed with its curing agent at a 5:1 mass ratio and the mixture was degassed. The pre-cured PDMS was then poured on the molds slowly and degassed again to remove the air bubbles trapped between the mold and the pre-cured PDMS. Molds were then placed in a 60°C convection oven for 90 minutes. After curing, PDMS was removed from the mold and holes were punched with a 16-gauge blunt tip needle (Cat: 5FVG5, Grainger, USA) for inlet and outlet connections. Later, patterned surface of PDMS was cleaned with tape and irreversibly bonded to a 24 mm × 60 mm #2 pre-cleaned cover glass (Warner Instruments, USA) by oxygen plasma bonding (Harrick Plasma Cleaner, 30 seconds, 18 W). The assembly was placed in a convection oven at 90°C for 2 hours to facilitate covalent bonding between the glass and PDMS. Microchannels were stored at room temperature overnight before use and discarded after the experiment.

### Scanning Electron Microscopy (SEM)

Prior to SEM imaging, PDMS parts were cleaned with tape and filtered nitrogen stream. For the cross-section imaging, the PDMS was gently cut using a double edge carbon steel razor blade (Cat: 71933-50, Electron Microscopy Sciences, USA), placed onto double-sided tape that had been adhered to a glass slide, and sputter coated with an approximately 50 Å gold layer. SEM images were acquired using a Jeol JSM 6400 scanning electron microscope, with accelerating voltages ranging from 10 kV to 25 kV and magnification ranging from 40x to 4000x.

### Numerical Flow Characterization

Computational Fluid Dynamics (CFD) simulations were performed using 3D COMSOL (Comsol Inc., Sweden) model to numerically predict the wall shear stress magnitudes at every location within the microfluidic device. Using the laminar flow assumption with Newtonian perfusion media and no-slip condition at the microchannel walls, the Navier-Stokes equation was solved for 2,431,573 elements that were created on the 3D microchannel model with physics-controlled fine-meshing (average mesh size: 20.19 µm). As a result, 3D velocity profiles were obtained across the microchannel and shear rate was calculated. The experimentally-measured dynamic viscosity value of the medium at 37°C was used in the simulation, and shear stress values for each element were calculated by multiplying viscosity and the shear rate using the linear proportionality property of Newtonian fluids.

### Microfluidic Perfusion Experiment Setup

All parts used in flow experiments that were in contact with the perfusion medium were autoclaved (121°C, 15 PSI, 30 minutes) and the flow circuit was assembled in a sterile cell culture hood. First, peristaltic pump tubing with stoppers (Cat: EW-06447-13, Cole Parmer, USA) was installed in a peristaltic pump (7524-50 Master flex L/S with 7518-10 pump head, Cole Parmer, USA) and connected to the 1/32” ID 3350 Tygon tubing (Cat: 025871A, Fisher Scientific, USA) with a 1/16” straight polypropylene barbed fitting (Cat: 5121K191, McMaster-Carr, USA) at both ends. The upstream flow tubing was connected to a 50-mL media reservoir and the downstream flow tubing was connected to a custom flow dampener. The outlet of the flow dampener was then connected to the inlet of the bubble trap (Diba Omnifit PEEK with 10 µm PTFE filter, Cole Parmer, USA) using Tygon tubing and quick-turn polypropylene tube couplings (Cats: 51525K291; 51525K141, McMaster-Carr, USA). After these connections were made, the flow circuit was primed with culture media at 3.6 mL/min. The perfusion circuit was visually inspected for trapped air bubbles before connecting the microfluidic device. Finally, the outlet of the bubble trap was connected to the inlet of the microfluidic system by rigid polytetrafluoroethylene (PTFE) tubing (Cat: 5239K23, McMaster-Carr, USA) with a flow rate of 0.2 ml/min. The outlet of the microfluidic device was connected back to the reservoir unit using PTFE tubing and Tygon tubing to complete the assembly of the closed-circuit perfusion system. The assembled perfusion setup was placed in an incubator and the flow rate of the pump was increased to the desired flow rate after 30 minutes.

For live imaging, the perfusion system was set up in the same way but instead of placing the entire system in a cell culture incubator, the microfluidic device was placed in a stage top environmental chamber (STXF-WSKMX-SET, Tokai Hit, Japan) at 37°C and 5% CO_2_. The flow reservoir and pulsation dampener was kept at 37°C near the microscope stage. For the control experiments with live imaging, the cell culture media in the microchannels was slowly replenished using a syringe pump at the flow rate of 0.4 µL/min such that the maximum shear stress in the microchannels would not exceed 0.1 dyne/cm^2^.

### Perfusion Media Viscosity Measurement

The dynamic viscosity of the cell culture media was measured with AR2000 plate-cone rheometer (TA Instruments, USA). An 80 mm diameter cone with the cone angle of 1° was set to 50 µm away from the plate, and data were collected at different steady state shear rate values ranging from 500 to 2500 s^−1^. In this way, the Newtonian behavior of the cell culture medium was confirmed and the average dynamic viscosity values across a large shear rate range were calculated. All measurements were taken at 37°C (± 0.1°C). As a control, viscosity of filtered deionized (DI) water was measured with the same settings.

### Experimental Flow Characterization

The flow system was set up as described above (without cells), with the microfluidic device placed on the stage of a spinning disk confocal (Yokagawa CSU-X1, Hamamatsu X2 EMCCD mounted on a Leica DMI-6000) and perfused with fluorescent beads (0.2 µm in diameter, Cat: F8811, Thermo Fisher Scientific, USA) in phosphate-buffered saline (PBS, Thermo Fisher Scientific, USA) at 0.5 ml/min. After 15 minutes, images were collected with an exposure time of 30 milliseconds. In the low uniform shear stress region, the entire Z stack was collected and analyzed to plot the flow profile across the microchannel height. For the flow visualization experiments, polystyrene green fluorescence beads (1 µm in diameter, Thermo Fisher Scientific, USA) at 1% w/v were perfused into the microfluidic device and the videos of the streaklines of the particles were acquired using an Olympus IX83 inverted fluorescence microscope.

### Primary Endothelial Cell Culture

Pooled HUVECs (C-12203, Promocell, Heidelberg, Germany) were cultured at 37°C under 5% CO_2_ in a sterile humidified incubator in complete growth medium (Endothelial Cell Basal Medium-2 C-22211, supplemented with Endothelial Cell Growth Medium 2 Supplement Pack C-39211, Promocell) and 1X Antibiotic-Antimycotic (Cat: 15240112, Thermo Fisher Scientific, USA). Upon confluency, HUVECs were rinsed with PBS, treated with 0.05% trypsin-EDTA (Cat: 25300062, Thermo Fisher Scientific, USA) for 5 minutes at room temperature, centrifuged at 300 × g for 4 minutes, and gently resuspended in fresh culture medium for a maximum of six passages.

### Microfluidic Cell Seeding and Culture

Microfluidic channels were autoclaved (121°C, 15 PSI, 30 minutes) and filled with sterile ethanol. Then, ethanol was replaced with sterile DI water and channels were coated with 50 µg/mL fibronectin (Cat: 10838039001, Sigma Aldrich, USA) at 37°C for 1 hour in a cell culture incubator. HUVECs were suspended at 3 million cells per ml in complete growth media, and 40 µl of this suspension was slowly injected into the fibronectin-coated microfluidic device from the inlet hole using a P200 micropipettor. The excess media around the outlet was immediately cleaned with sterile wipes and the device was placed in a petri dish and kept in the incubator for 30 minutes. Once HUVECs began to adhere to the fibronectin-coated glass slide, 1 ml culture media was added on top of the inlet and outlet ports for overnight culture. Cells were cultured in microchannels for at least 16 hours to confluency to ensure that a uniform HUVEC monolayer was established (Supplementary Figure 5) before using them in the flow experiments. For static control experiments, the same seeding procedure was performed and the media was replaced daily.

### Cell Fixing, Staining, and Endpoint Imaging

For all cell fixing and staining procedures, reagents were pipetted into microfluidic channels three times to ensure equal reagent delivery to all cells in different regions of the device. Immediately after the perfusion experiments, cells were fixed with 4% pre-heated paraformaldehyde (PFA) for 15 minutes in the 37°C incubator and stored in PBS at 4°C until immunofluorescence (IF) staining. For IF, all steps were performed at room temperature, within the microchannel. Cells were permeabilized with 0.1% (v/v) Triton X-100 (Cat: T8787, Sigma Aldrich, USA) in PBS (PBST) for 15 minutes and blocked with 10% (v/v) goat serum (Cat: G9023, Sigma Aldrich, USA) in PBST for 30 minutes. Golgi were labeled using mouse anti-GM130 (1:200 dilution, Cat: 610823, BD Biosciences, USA) and goat anti-mouse IgG-rhodamine RedX (1:500 dilution, Cat: 115-295-003, Jackson ImmunoResearch Laboratories, USA). All antibodies were incubated for 1 hour at room temperature. Samples were then rinsed with PBST and nuclei were stained for 10 minutes with 5 µg/ml Hoescht 33342 (Cat: H3570, Thermo Fisher, USA). Imaging was performed using a Nikon A1 confocal microscope on an inverted Nikon TiE platform (Nikon Instruments, Japan).

### Live Imaging

HUVECs at ∼50% confluency were transduced with CellLight Golgi-GFP (BacMam 2.0, Thermo Fisher, USA), 2 µl per 10,000 cells, resulting in 70% trandsuction efficiency when assayed at 16 hours. Transduced cells were then seeded into the microchannels as described above and the nuclei were stained with 1 µg/ml Hoescht 33342 for 20 minutes before applying flow. Fluorescent and DIC images were acquired every 15 minutes for 6 hours. A modified Nikon Ti-E (Nikon instruments, Japan) was used for these studies. The brightfield illumination system used differential interference contrast with 505 nm transmitted light illumination from a diode illuminator (ScopeLED, USA) at 15 ms exposure time. The fluorescence images used a Lumencor Spectra X light source (Lumencor, USA). Images were collected with a Teledyne-Photometrics 95B camera (Teledyne-Photometrics, USA). Each image was composed of a field of 28 × 7 3-color images (GFP, Hoescht, DIC) collected sequentially and montaged to create the overview of the entire microfluidic chip. To maximize speed the images were collected using a high speed 5 position dichroic changer (50 ms change time) and high-speed filter wheel (10 ms change time) (FLI instruments, USA). Focus across the entire field was maintained using the Nikon perfect focus mechanism. Images were collected with a 10x 0.5NA plan apochromatic objective and montaged using Nikon NIS Elements software (5.1.01).

### Automated Analysis of Golgi and Nucleus

Analysis of the Golgi and nucleus morphology from fluorescence endpoint images was automated using a custom-written macro for Fiji ^63^. Nuclei and Golgi were segmented independently and paired based on the distance between them. Among these pairs, if the center of the Golgi was within the nucleus region, the cell was designated as unpolarized. If the center of the Golgi was outside the nucleus region, the cell was designated as polarized and further characterized according to its orientation with respect to the flow direction. Cells were binned into four categories: upstream-oriented (Golgi-nucleus vector aligned against the direction of flow), downstream-oriented (Golgi-nucleus vector aligned with the direction of flow), right-oriented, or left-oriented. The macro returned the number of cells analyzed, percentage of the cells in each polarization category, the average distance between the centers of the Golgi and nucleus, the average area of the Golgi and nucleus, as well as parameters needed for subsequent statistical analyses. This macro allowed objective analysis of thousands of cells within a set of images in a few minutes. A flow chart detailing this macro can be found in Supplementary Figure 6.

### Statistical Methods

Every experiment was repeated at least 3 times and the quantitative parameters were calculated for each experiment independently. A two-tailed unpaired student’s *t*-test was applied for statistical comparison of the data from two different conditions. The cases where p > 0.05 were considered not significant (ns). Experiments with more than two conditions were analyzed using one-way ANOVA followed by *post hoc* two-tailed unpaired student’s *t*-test with Bonferroni correction for pairwise comparison. Categorical data were analyzed by chi-square independence test followed by post hoc adjusted residual test with Bonferroni correction for pairwise comparison.

## Acknowledgements

We would like to thank Derrick Amoabeng and Sachin Velankar for their help with viscosity measurement and University of Pittsburgh Center for Biological Imaging (CBI) staff for live imaging. We also appreciate William Okech and Arulselvi Anbalagan (Human Genetics, University of Pittsburgh) for stimulating discussions throughout the study. This work was supported by the National Institutes of Health (R01HL136566).

## Author Contributions

B.L.R. and L.A.D. conceived the initial research idea. U.M.S. designed and fabricated the microfluidic system. U.M.S. and Y.W.C. conducted the experiments. All authors contributed to the interpretation of the results. U.M.S wrote the manuscript with support from all authors. All authors read and approved the final version of the manuscript.

## Data availability

The relevant data are available from the authors upon reasonable request.

