## Supplementary Materials for "Endothelial Cell Polarization and Orientation to Flow in a Novel Microfluidic Multimodal Shear Stress Generator"

### Supplementary Information:

The solution of the Navier-Stokes equations for the steady-state laminar flow of an incompressible Newtonian fluid between two infinitely-long parallel plates leads to a well-known flow regime known as Hele-Shaw flow. Since the microfluidic channels generally have low aspect ratio (i.e. height << width), the hydrodynamic characterization of the flow can be initially simplified with this parallel plate assumption. The parabolic flow profile of Hele-Shaw flow demonstrates a parabolic velocity distribution across the height of the gap in the following form

$$v(z) = \frac{6}{h^2} z(h - z)\underline{v} \qquad \text{(S1)}$$

where $\underline{v}$ is the mean velocity, $h$ is the height of the gap, and $z$ is the coordinate along the height of the gap starting as zero from the bottom surface of the microchannel. In this way, the wall shear stress acting on the cells in the microchannel regions with uniform width can be approximated as

$$\tau_w = \frac{6\mu Q}{wh^2} \qquad \text{(S2)}$$

where $\mu$ is the dynamic viscosity of the fluid, $Q$ is the volumetric flow rate, and $w$ is the width and $h$ is the height of the microchannel. Microchannel regions with changing width are expected to have linear stream-wise shear stress gradients (i.e., along the $x$ axis) in order to expose cells consistently to the same gradient slope in the given region. The linear shear gradient can be expressed in the form of

$$\tau_w(x) = ax + b \qquad \text{(S3)}$$

where $a$ and $b$ can be found by solving this equation at the boundaries of the linear shear stress gradient regions. Using the Hele-Shaw flow assumption, the wall shear stress at the beginning of the linear shear stress gradient region can be calculated as

$$\tau_w(x)_{[x=0]} = b = \frac{6\mu Q}{w_i h^2} \qquad \text{(S4)}$$

where $w_i$ is the width of the microchannel at the beginning of the linear shear stress gradient region at $x = 0$ (Supplementary Figure 1). Using the same Hele-Shaw flow assumption, the wall shear stress at the end of the linear shear stress gradient region can be similarly found as

$$\tau_w(x)_{[x=L]} = aL + \frac{6\mu Q}{w_i h^2} = \frac{6\mu Q}{w_f h^2} \tag{S5}$$

where $L$ is the length of the region and $w_f$ is the final width of the linear shear stress gradient region at $x = L$. From this equation $a$ can be found as

$$a = \frac{6\mu Q}{h^2}\frac{(w_i - w_f)}{L w_i w_f} \tag{S6}$$

which also represents the slope of the linear wall shear stress gradient generated in $x$-direction. Using these parameters, the magnitude of this wall shear stress at the given point along the $x$ axis can then be written as

$$\tau_w(x) = \frac{6\mu Q}{h^2}\frac{(w_i - w_f)}{L w_i w_f}x + \frac{6\mu Q}{w_i h^2} \tag{S7}$$

The formula for the function that describes the microchannel width can be derived from this equation as

$$w(x) = \frac{L w_i w_f}{(w_i - w_f)x + L w_f} \tag{S8}$$

Since there are two symmetrically curved sidewalls at each side of the microchannel that defines the width of the channel, the formula needed for plotting each sidewall becomes

$$c(x) = \frac{L w_i w_f}{2((w_i - w_f)x + L w_f)} \tag{S9}$$

which can be used to model the microchannel regions with linear shear stress gradient in a CAD software.

### Supplementary Figures:

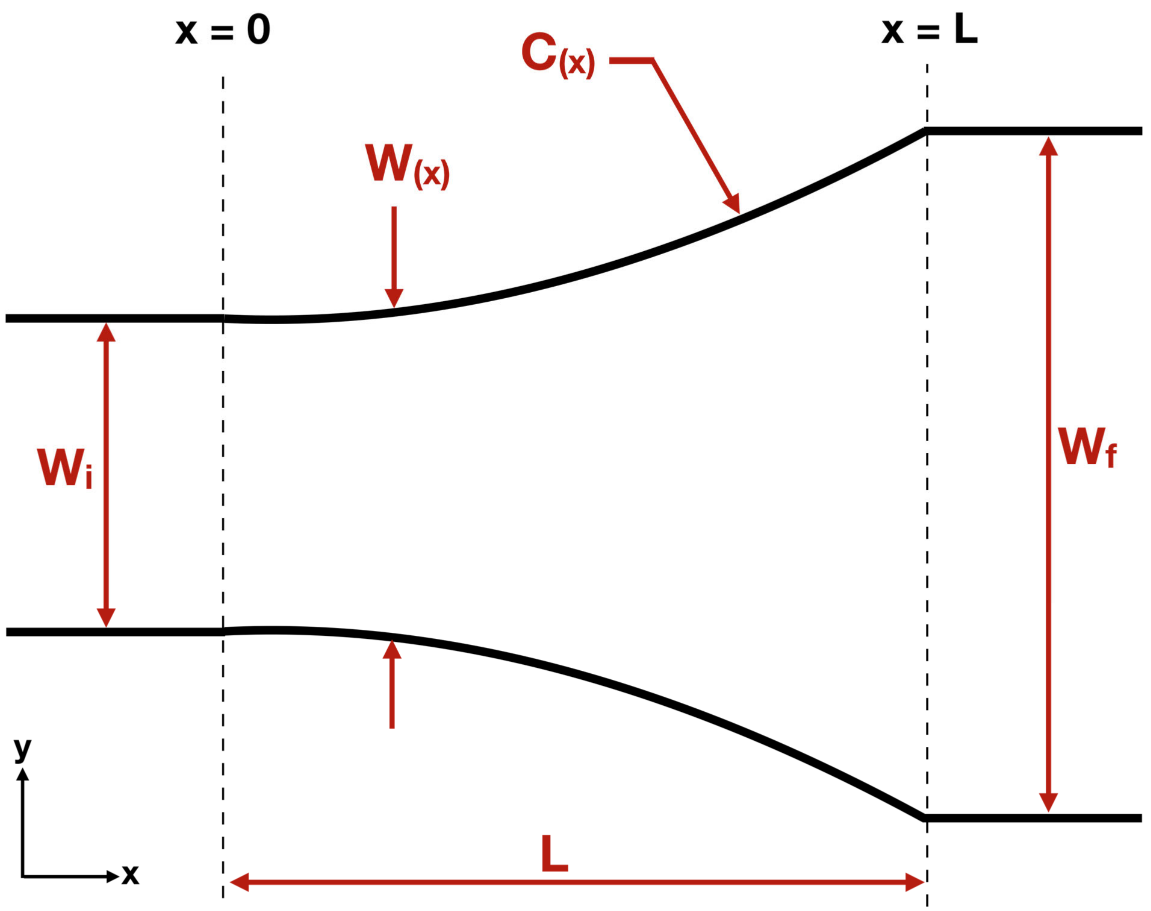

**Supplementary Figure 1: Annotations of the parameters used for analytical characterization of the streamwise wall shear stress gradients in the microfluidic chip.**

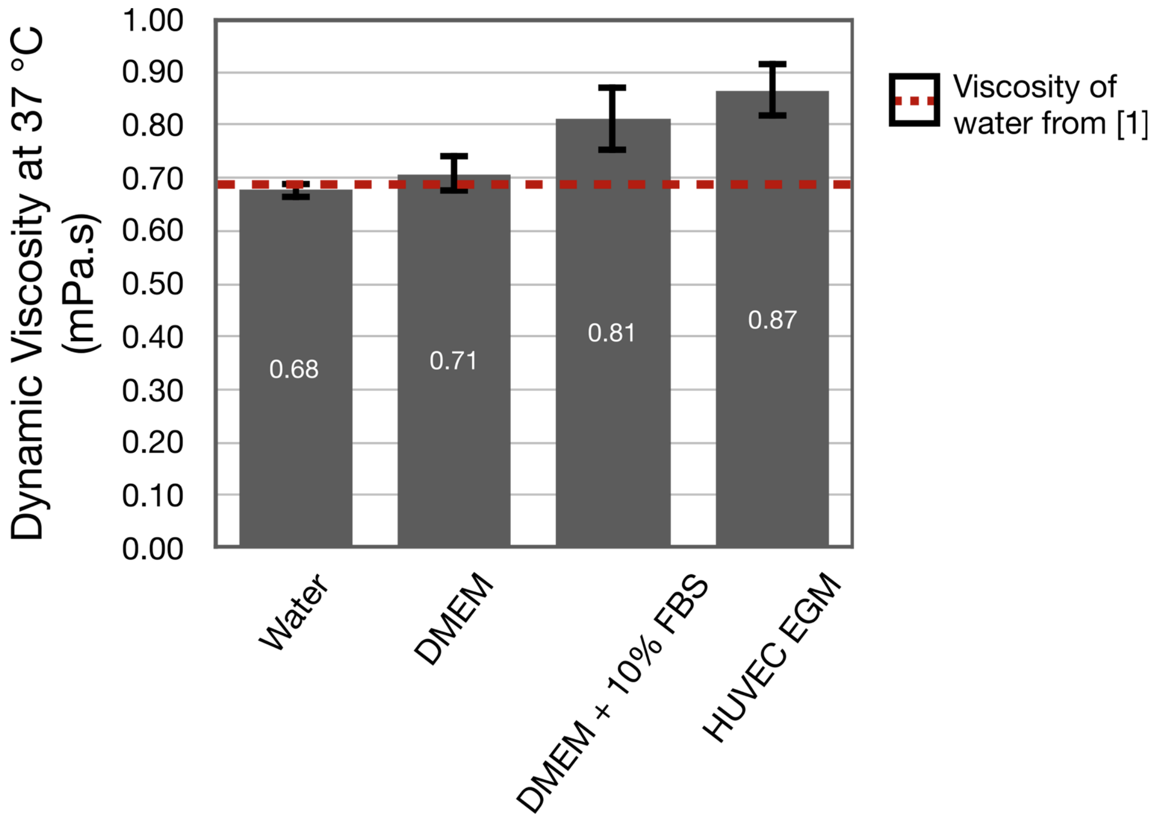

**Supplementary Figure 2: Dynamic viscosity values of various cell culture media at 37°C.** The average value of the water viscosity was 2.89% different than the value in the literature. Addition of 10% FBS (Fetal Bovine Serum) into DMEM (Dulbecco's Modified Eagle Medium) increased dynamic viscosity of the media. The complete HUVEC EGM (Human Umbilical Cord Vascular Endothelial Cell Growth Media) that was used throughout this study had even higher viscosity. Data are presented as mean ± SD.

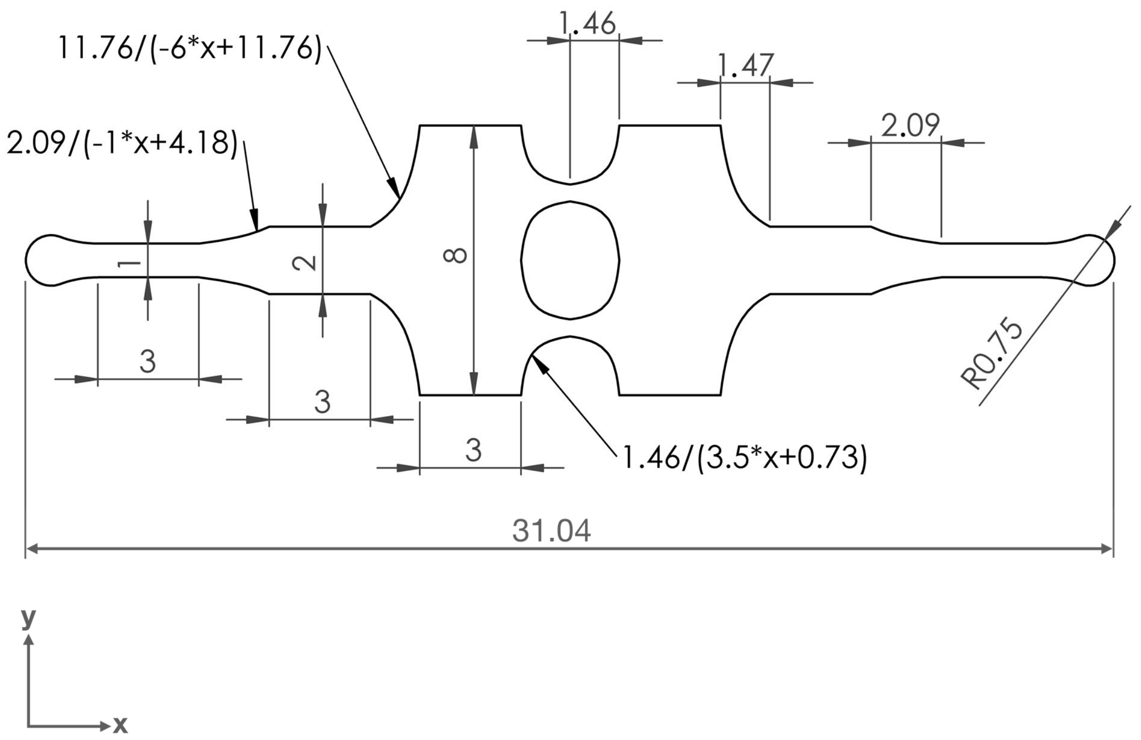

**Supplementary Figure 3: The dimensions of the microfluidic chip used in the experiments.** The equations describing the curved sidewalls were generated using Equation 6 in the manuscript. The height of the different microchannels regions is constant and equal to 200 µm. The microfluidic chip is symmetric with respect to x- and y-axes. All units are in mm.

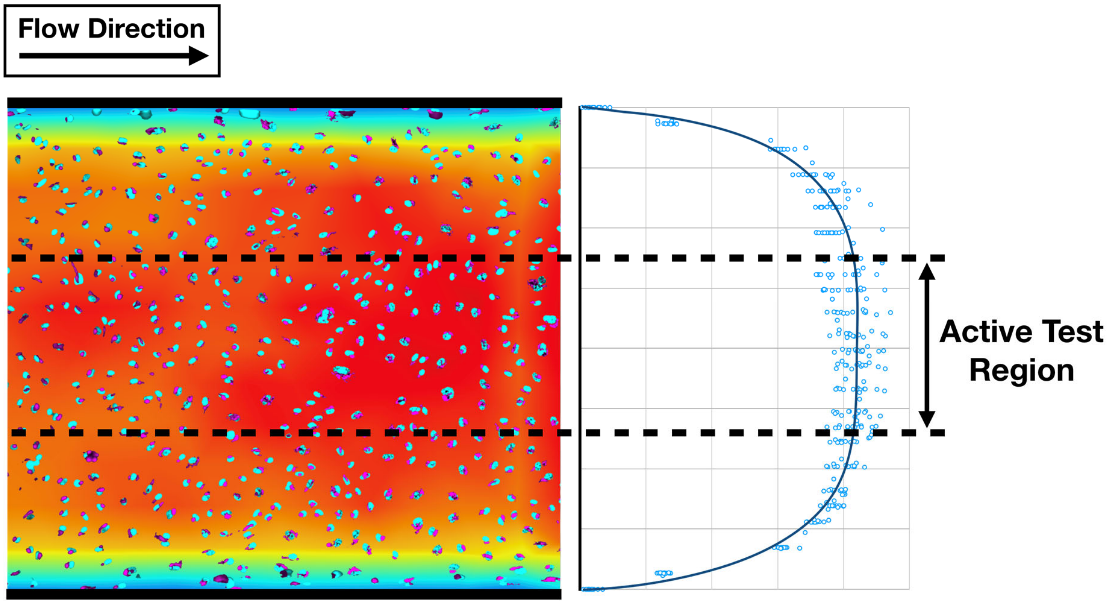

**Supplementary Figure 4: Computational Fluid Dynamics (CFD) simulation results showed the distribution of the shear stress across the bottom surface of the microchannel where the endothelial cells were seeded.** Fluorescent images of Golgi and nuclei of the human umbilical vein endothelial cells (HUVECs) overlaid with the shear stress map. Due to the sidewalls, the shear stress exerted on the cells was lower at either side of the microchannel. Therefore, only the HUVECs in the central region of the microchannel (shown between two parallel dashed lines) were analyzed.

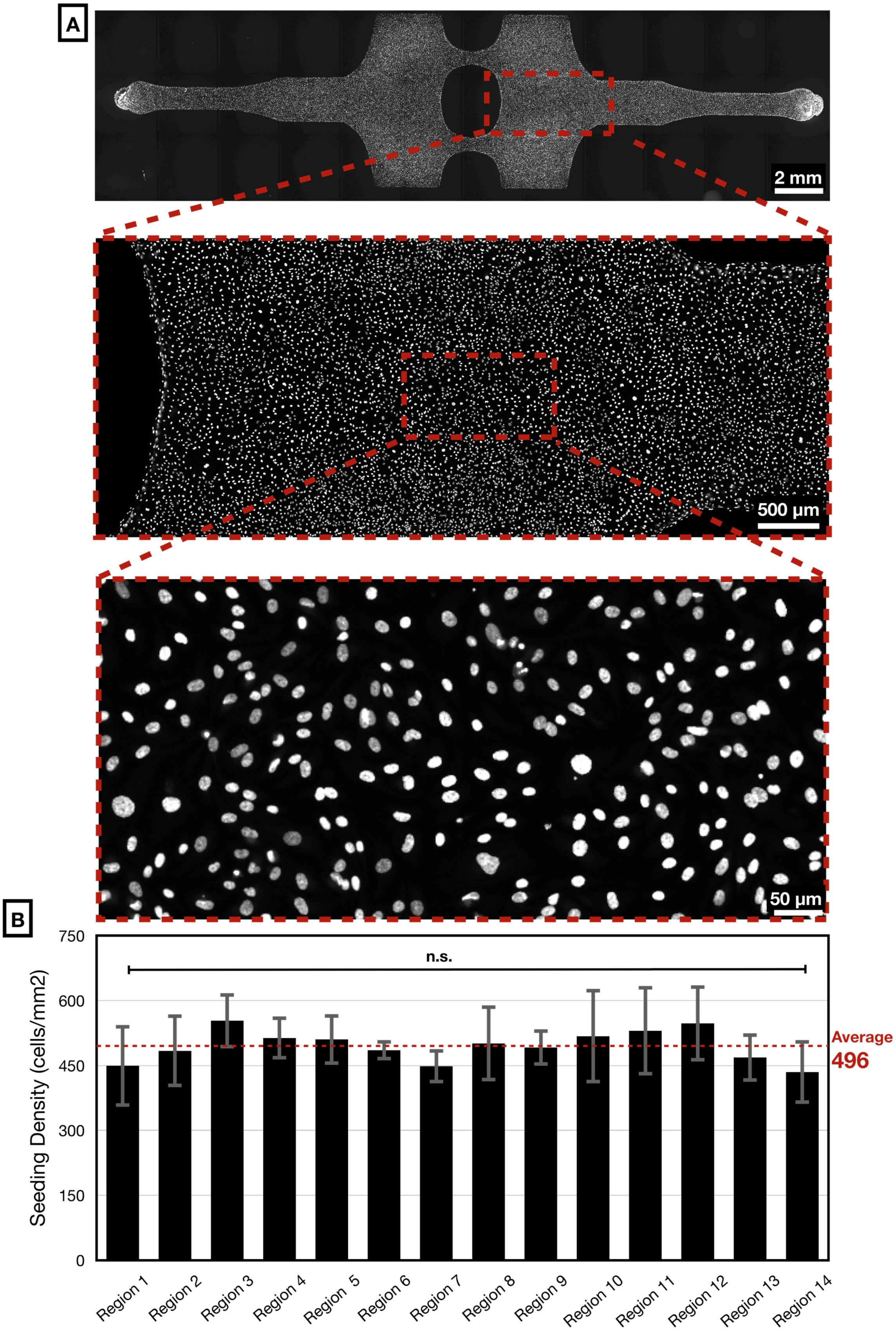
A
2 mm
500 µm
50 µm
B
n.s.
Average
496
Seeding Density (cells/mm2)
750
600
450
300
150
0
Region 1
Region 2
Region 3
Region 4
Region 5
Region 6
Region 7
Region 8
Region 9
Region 10
Region 11
Region 12
Region 13
Region 14

**Supplementary Figure 5: Uniform endothelial cell monolayer across the entire microfluidic chip**.

a) Nuclei were stained with Hoescht 33342 and imaged with an epifluorescence microscope. Images show the distribution of the cells across the microfluidic chip with increasing magnification.

b) Quantitative analysis of the cell seeding density showed the number of HUVECs per $mm^2$ of each region in the microfluidic chip, with the red dashed line showing the average of all the regions. The number of cells increased as the microchannels got wider. This would be expected as the cells can more easily attached to surface fibronectin when they slowed down in large microchannel regions. However, this change was not statistically significant ($p=0.178$ by one-way ANOVA). Data are presented as mean ± SD.

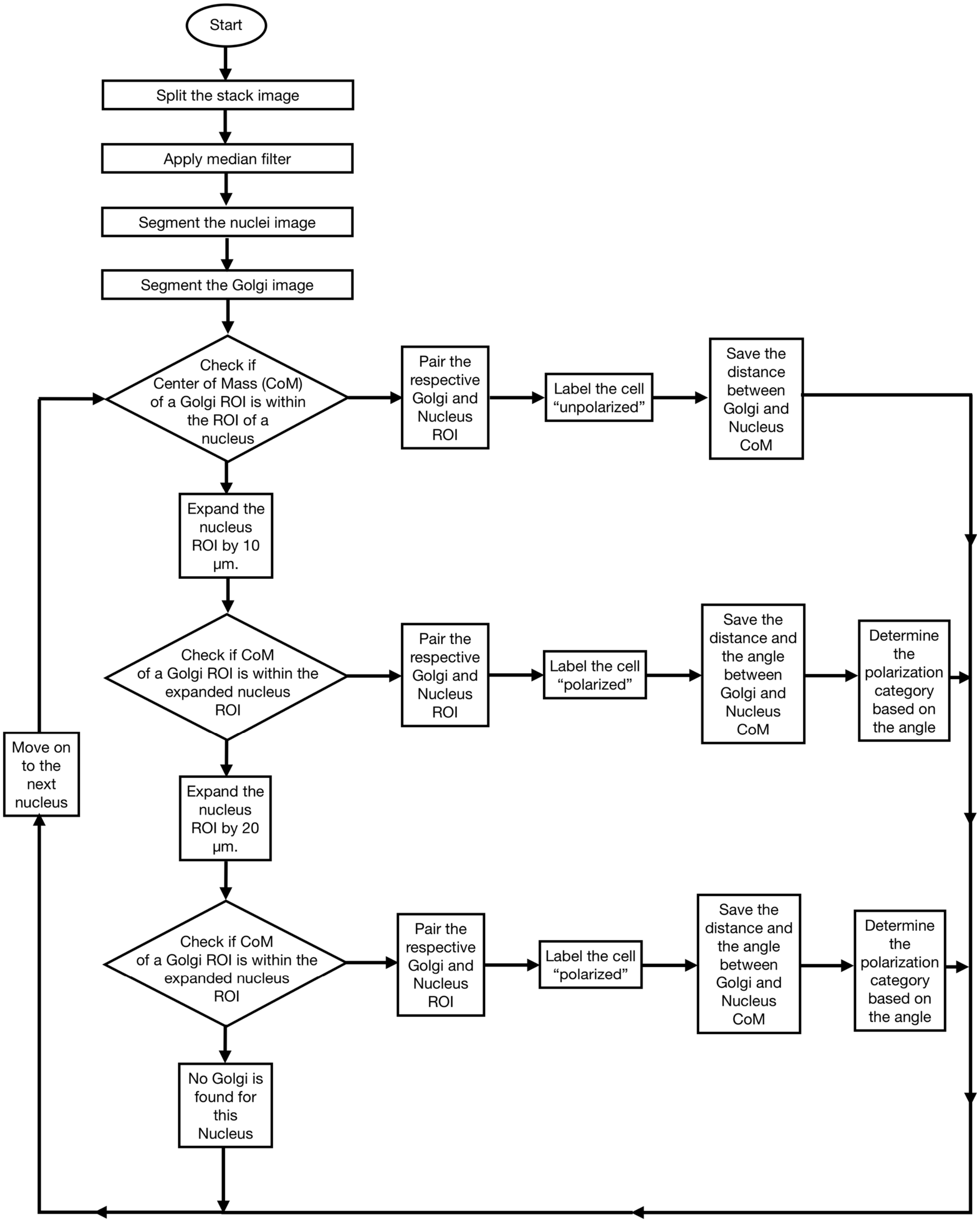

**Supplementary Figure 6: Flowchart of the macro for automated Golgi-nucleus polarization and orientation analyses.** After the preprocessing for lowering the noise, the Golgi and nuclei are segmented and paired. The cells where the center of mass (CoM) of Golgi are outside the nucleus are designated as polarized. The polarized cells are binned into four categories based on the angle of the vector connecting the center of the nucleus and Golgi. Within a 45° interval, if the Golgi is upstream of the nucleus, the cell is designated as upstream polarized. If the Golgi is downstream of the nucleus, the cell is designated as downstream polarized. If the Golgi is to the side of the nucleus, the cell is designated as right or left polarized. The macro produces a polarization map image where all nuclei are labelled with different colors based on their polarization category.

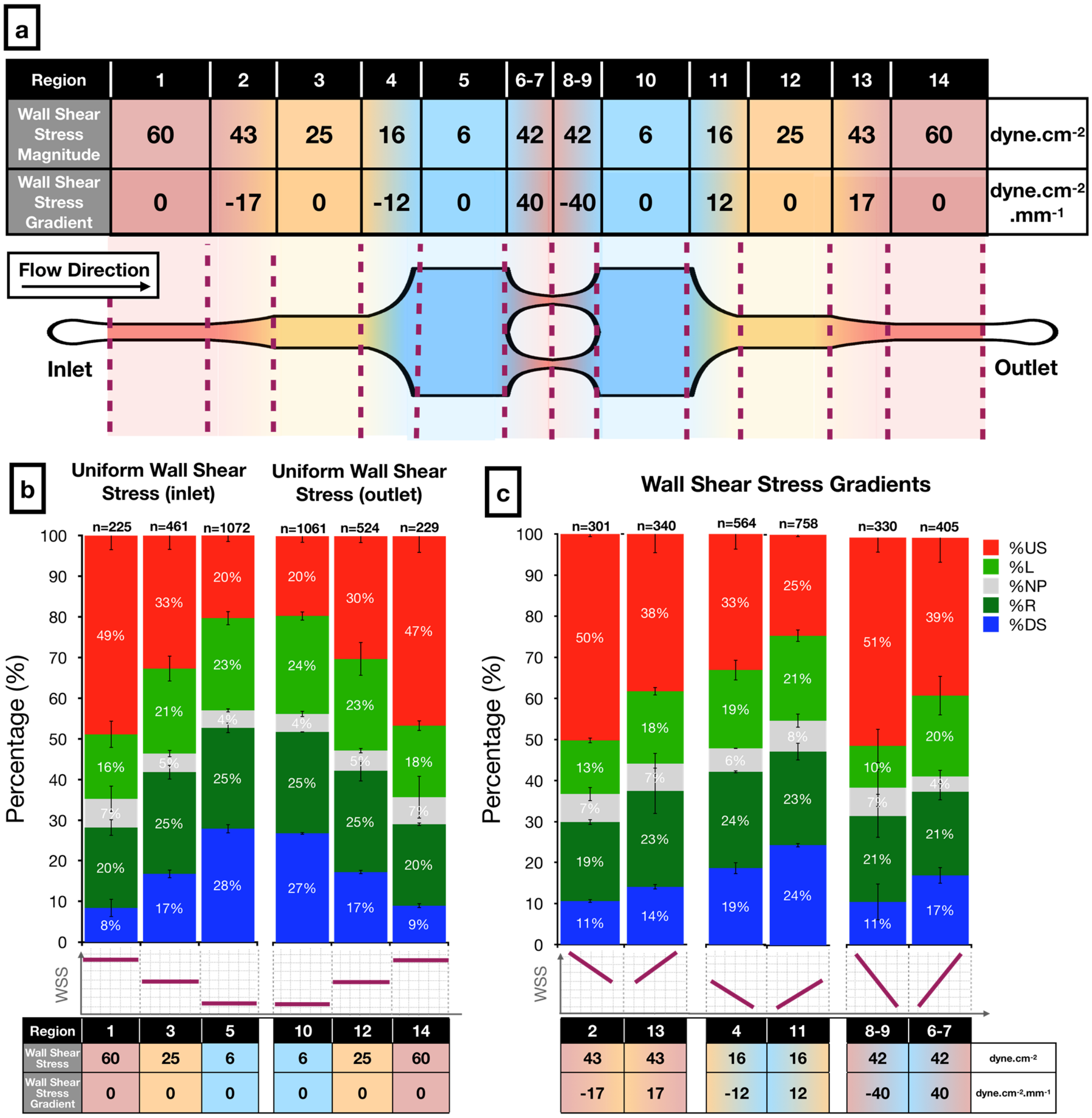

**Supplementary Figure 7: Golgi-nucleus polarization and orientation in a microfluidic chip with different dimensions that create higher magnitudes of wall shear stress (WSS) and steeper wall shear stress gradients (WSSG).** Confluent human umbilical vein endothelial cells (HUVECs) were exposed to different shear stress modalities for 24 hours.

a) Schematic and table showing the magnitudes and slopes of the WSS and WSSG, respectively, across the microfluidic chip.

b) Stacked bar graphs showing Golgi-nucleus orientation patterns of HUVECs in response to different levels of uniform WSS.

c) Stacked bar graphs showed the Golgi-nucleus polarization patterns of HUVECs that were exposed to different WSSGs.

In b and c, Golgi-nucleus orientation is indicated as upstream (red), downstream (blue), right (dark green), left (light green), or nonpolarized (gray). n = number of cells analyzed in each region. Data from a single experiment; mean ± SEM.

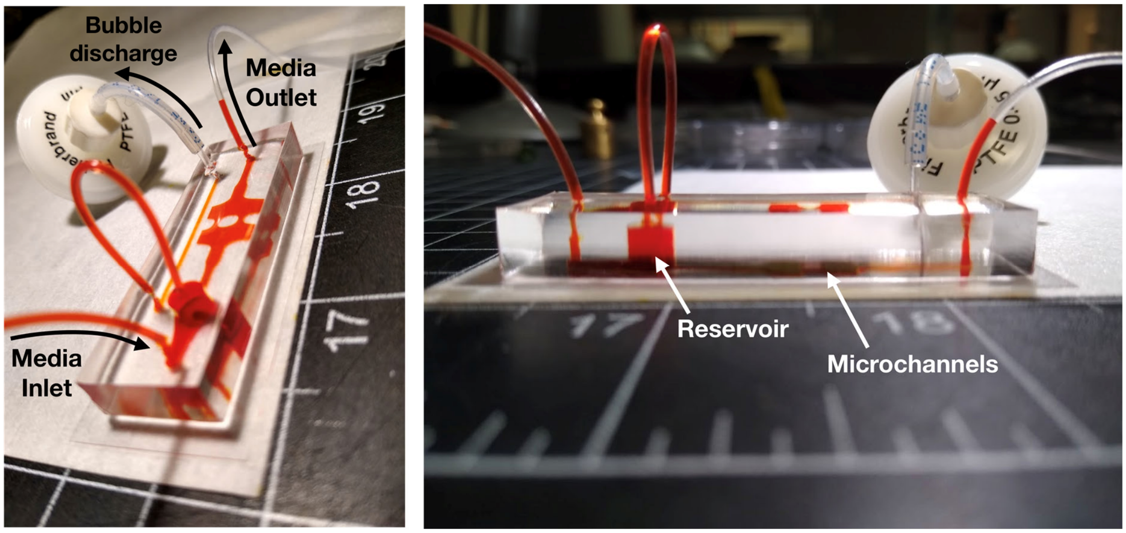

**Supplementary Figure 8: The on-chip integration of the bubble trap**. Photos show the microfluidic chip with the integrated bubble trap for decreasing the "lab-around-the-chip". All the microchannels were filled with red food dye to facilitate visualization. The 2 mm diameter reservoir upstream of the microchannels was connected to a narrow bubble discharge channel that was plugged with a syringe filter to maintain sterility inside the microchannel. As the bubbles entered to the microfluidic chip through the inlet tubing, they rose and trapped in the reservoir region due to their buoyancy. Over time, these bubbles were pushed into the bubble discharge channel because of the positive pressure inside the microfluidic chip and they left the system through the membrane filter. In this way, the microchannels could be maintained bubble-free without using an external debubbler.

### Supplementary Tables

**a**

Static Control: Categorical Golgi-nucleus Polarization-Orientation Distribution Comparison

| Microchannel Regions | P-values |
|---|---|
| Uniform WSS (Figure 4b) | |
| 1 vs 3 | 0.684 |
| 3 vs 5 | 0.937 |
| 10 vs 12 | 0.424 |
| 12 vs 14 | 0.736 |
| WSSG (Figure 5a-c) | |
| 2 vs 13 | 0.364 |
| 4 vs 11 | 0.998 |
| 6 vs 8 | 0.764 |
| 7 vs 9 | 0.461 |

**b**

24H Flow: Upstream Orientation Comparison

| Microchannel Regions | P-values |
|---|---|
| Uniforms WSS (Figure 4b) | |
| 1 vs 3 | 1.02E-05 |
| 3 vs 5 | 2.00E-06 |
| 10 vs 12 | 3.34E-02 |
| 12 vs 14 | 1.14E-05 |
| WSSG (Figure 5a-c) | |
| 2 vs 13 | 5.65E-03 |
| 4 vs 11 | 2.25E-04 |
| 6 vs 8 | 1.16E-04 |
| 7 vs 9 | 7.87E-05 |

**c**

Static Control vs. 24H Flow: Categorical Golgi-nucleus Polarization-Orientation Distribution Comparison

| Microchannel Regions | P-values |
|---|---|
| 1 | <1.00E-16 |
| 2 | <1.00E-16 |
| 3 | <1.00E-16 |
| 4 | <1.00E-16 |
| 5 | 3.35E-12 |
| 6 | <1.00E-16 |
| 7 | <1.00E-16 |
| 8 | <1.00E-16 |
| 9 | <1.00E-16 |
| 10 | 4.77E-09 |
| 11 | 1.01E-13 |
| 12 | <1.00E-16 |
| 13 | <1.00E-16 |
| 14 | <1.00E-16 |

**Supplementary Table 1: The list of exact p-values for comparison of the polarization and orientation patterns in response to 24 hours-long exposure to different shear stress modalities.**

a) Comparison of the polarization and orientation responses of the HUVECs from different microchannel regions in static control experiments.

b) Comparison of the upstream orientation among HUVECs subjected to different shear stress modalities.

c) Comparison of the polarization and orientation responses of the HUVECs in flow and static control experiments. P-values were obtained by chi-square independence test followed by post hoc test with adjusted residuals using Bonferroni correction for pairwise comparison. (WSS: Wall shear stress; WSSG: Wall shear stress gradient. See Figure 4 and 5 for the numerical values of WSS and WSSG in each microchannel region).

Uniform WSS vs WSSG: Categorical Golgi-nucleus Polarization-Orientation Distribution Comparison

| Microchannel Regions | P-values |
|---|---|
| High Uniform WSS vs. Steep Negative WSSG | |
| 1 vs 8 | 4.51E-02 |
| 1 vs 9 | 8.65E-01 |
| 14 vs 8 | 7.15E-01 |
| 14 vs 9 | 5.86E-01 |
| Low Uniform WSS vs. Shallow Positive WSSG | |
| 5 vs 11 | 5.98E-01 |
| 10 vs 11 | 7.14E-01 |

**Supplementary Table 2: The list of exact p-values for comparison of the polarization and orientation patterns in response to uniform WSS and WSSGs.**

P-values were obtained by chi-square independence test. (WSS: Wall shear stress; WSSG: Wall shear stress gradient. See Figure 4 and 5 for the numerical values of WSS and WSSG in each microchannel region).

**a**

P-values: Static Control vs. Shear Stress Responses Comparison

| Microchannel Regions | Nucleus Area | Golgi Area | Golgi-Nucleus Distance |
|---|---|---|---|
| Uniforms WSS (Figure 7) | | | |
| 1&14 | 6.32E-03 | 3.47E-03 | 9.62E-05 |
| 3&12 | 1.37E-02 | 3.97E-03 | 1.86E-04 |
| 5&10 | 1.79E-02 | 9.57E-04 | 1.18E-05 |
| WSSG | | | |
| 2 | 4.15E-02 | 1.95E-04 | 2.83E-03 |
| 13 | 1.70E-02 | 7.74E-03 | 1.80E-05 |
| 4 | 1.66E-02 | 8.43E-03 | 6.45E-04 |
| 11 | 1.66E-03 | 7.18E-03 | 2.06E-04 |
| 8&9 | 6.11E-03 | 6.30E-03 | 1.05E-03 |
| 6&7 | 1.03E-03 | 5.88E-04 | 1.60E-04 |

**b**

P-values: Shear Stress Responses Comparison

| Microchannel Regions | Nucleus Area | Golgi Area | Golgi-Nucleus Distance |
|---|---|---|---|
| Uniforms WSS (Figure 7) | | | |
| 1&14 vs 3&12 vs 5&10 | 7.60E-01 | 5.66E-01 | 3.24E-01 |
| WSSG | | | |
| 2 vs 13 | 9.30E-01 | 8.28E-01 | 2.02E-01 |
| 4 vs 11 | 8.15E-01 | 7.92E-01 | 2.32E-01 |
| 8&9 vs 6&7 | 9.89E-01 | 2.83E-01 | 7.12E-01 |

**Supplementary Table 3: The list of exact p-values for comparison of Golgi area, nucleus area, and Golgi-nucleus distance after 24 hours of shear stress.**

a) Comparison of the shear stress responses with static control responses for the same microchannel regions.

b) Comparison of responses to different shear stress modalities. P values were obtained by one-way ANOVA followed by post hoc two-tailed unpaired student's t-test with Bonferroni correction for pairwise comparison. (WSS: Wall shear stress; WSSG: Wall shear stress gradient. See Figure 4 and 5 for the values of WSS and WSSG in each microchannel region).
